# Muscle-secreted neurturin couples myofiber oxidative metabolism and slow motor neuron identity

**DOI:** 10.1101/2021.03.31.437883

**Authors:** Jorge C. Correia, Yildiz Kelahmetoglu, Paulo R. Jannig, Christoph Schweingruber, Dasa Svaikovskaya, Liu Zhengye, Igor Cervenka, Mariana Oliveira, Jik Nijssen, Vicente Martínez-Redondo, Michael Stec, Naveen Khan, Johanna T. Lanner, Sandra Kleiner, Eva Hedlund, Jorge L. Ruas

**Affiliations:** Molecular and Cellular Exercise Physiology, Department of Physiology and Pharmacology, Biomedicum. Karolinska Institutet, 17165 Stockholm, Sweden; Department of Neuroscience, Biomedicum. Karolinska Institutet, 17165 Stockholm, Sweden; Molecular Muscle Physiology and Pathophysiology, Department of Physiology and Pharmacology, Biomedicum. Karolinska Institutet, 17165 Stockholm, Sweden; Regeneron Pharmaceuticals, Inc., Tarrytown, NY 10 591, USA

## Abstract

Endurance exercise promotes skeletal muscle vascularization, oxidative metabolism, fiber-type switching, and neuromuscular junction integrity. Importantly, the metabolic and contractile properties of the muscle fiber must be coupled to the identity of the innervating motor neuron (MN). Here, we show that muscle-derived neurturin (NRTN) acts on muscle fibers and MNs to couple their characteristics. Using a muscle-specific NRTN transgenic mouse (HSA-NRTN) and RNA-sequencing of MN somas, we observed that retrograde NRTN signaling promotes a shift towards a slow MN identity. In muscle, NRTN increased capillary density, oxidative capacity, and induced a transcriptional reprograming favoring fatty acid metabolism over glycolysis. This combination of effects on muscle and MNs, makes HSA-NRTN mice lean with remarkable exercise performance and motor coordination. Interestingly, HSA-NRTN mice largely recapitulate the phenotype of mice with muscle-specific expression of its upstream regulator PGC-1α1. This work identifies NRTN as a myokine that couples muscle oxidative capacity to slow MN identity.

**HIGHLIGHTS:**

- NRTN is a myokine induced by physical exercise.
- Muscle-derived NRTN promotes a slow motor neuron identity.
- Muscle-derived NRTN enhances muscle oxidative metabolism.
- NRTN improves systemic metabolism, exercise performance and motor coordination.

## INTRODUCTION

It is widely recognized that regular physical exercise greatly benefits physical and mental health and has preventive and therapeutic applications in medicine. Therefore, identifying molecules that mediate the beneficial effects of exercise continues to be the goal of many research efforts worldwide. However, exercise is a complex intervention with direct and indirect effects on a variety of organs. Thus, increasing performance through training depends on cell-autonomous adaptations to exercise but also on mechanisms of inter-organ crosstalk coordinated by exchange of different signals. Many of those signals are secreted by contracting muscle fibers and include proteins (myokines) and other humoral factors such as metabolites (Neufer et al., 2015; Whitham and Febbraio, 2016). Many of the skeletal muscle responses to exercise are integrated in broad transcriptional programs regulated by PGC-1 coactivators (namely PGC-1α1 and isoforms and PGC-1β; reviewed in Correia et al., 2015; Martínez-Redondo et al., 2015). In particular, PGC-1α1 (Puigserver et al., 1998) has been intensively studied due to the remarkable phenotype of the corresponding skeletal muscle-specific overexpressor mice (MCK-PGC-1α1 transgenics, Lin et al., 2002). MCK-PGC-1α1 mice have 2-6-fold higher PGC-1α1 levels in skeletal muscle (depending on the muscle type) and without any training display many of the well-known adaptations to endurance training. These include increased muscle angiogenesis, mitochondrial biogenesis, oxygen transport, energy efficiency, fatty acid oxidation capacity, and most importantly, exercise performance (Agudelo et al., 2019; Arany et al., 2008a; Lin et al., 2002). Interestingly, MCK-PGC-1α1 mice have also been shown to be resistant to developing stress-induced depression-like behavior through a mechanism of inter-organ crosstalk involving muscle, brain, and adipose tissue, also linked to systemic energy expenditure (Agudelo et al., 2014, 2018).

One important consequence of increasing PGC-1α1 in muscle is a switch towards slow-oxidative fibers, which contribute to the endurance phenotype (Lin et al., 2002). Strikingly, the fiber-type switch induced by ectopic expression of PGC-1α1 in the muscle fiber (but not in the motor neuron, MN), maintains motor unit homogeneity, even when the switch is induced postnatally (Arnold et al., 2014; Chakkalakal et al., 2010). The mechanism for this retrograde signaling from muscle to MN, that ensures the slow – slow coupling is maintained, has remained unknown. We have previously identified neurturin (NRTN) as a PGC-1α1-controled myokine that promoted neuromuscular junction (NMJ) formation in an in vitro microfluidics cell co-culture system (Mills et al., 2018). NRTN is a member of the glial cell line-derived neurotrophic factor (GDNF) family, involved in the maturation of neuromuscular synapses during development and regeneration following nerve injury (Baudet et al., 2008). Here, we show that ectopic *Nrtn* expression in mouse skeletal muscle is sufficient to induce a shift towards a slow MN identity. Surprisingly, mice with muscle-specific *Nrtn* expression (HSA-NRTN transgenics) display a phenotype that largely recapitulates that of the MCK-PGC-1α1 mice, including reduced adiposity and robustly increased muscle oxidative metabolism, angiogenesis, and exercise performance. In addition, both HSA-NRTN mice and C57bl/6 mice transduced with an AAV8-NRTN to promote NRTN secretion from the liver, show improved metabolic parameters and motor coordination. Together, these results identify NRTN as a PGC-1α1 effector myokine with high therapeutic potential, that coordinates effects on muscle fibers and innervating MNs.

## RESULTS

### Skeletal muscle-derived NRTN promotes morphological and transcriptional MN remodeling

Physical exercise activates the expression of various PGC-1 coactivators in skeletal muscle, many of which promote an endurance-like phenotype (Martínez-Redondo et al., 2015). Interrogation of publicly available skeletal muscle transcriptomics data revealed that *Nrtn* transcriptional activation is a common feature to various PGC-1 coactivators associated with endurance adaptations, namely PGC-1α1, PGC-1α-b and PGC-1 β, and to the estrogen-related receptor gamma (ERRγ), an important transcriptional partner to PGC-1 coactivators and potent inducer of oxidative metabolism in skeletal muscle (Hatazawa et al., 2015; Narkar et al., 2011; Pérez-Schindler et al., 2012; Yadav et al., 2014) (Figure S1A). Conversely, skeletal muscle *Nrtn* expression is reduced in mice with genetic deletion of PGC-1α (Figure S1A). In line with the well-established role of PGC-1 coactivators in the physiological adaptation to physical exercise (Correia et al., 2015), *Nrtn* expression was elevated in mouse skeletal muscle after 8 weeks of voluntary wheel running (Figure 1A). Likewise, we observed a robust activation of *NRTN* expression in human vastus lateralis muscle after a single session of high-intensity sprint cycling (Schlittler et al., 2019) (Figure 1B). Consistent with these results, *Nrtn* expression in skeletal muscle correlates positively with the improvement of VO2 max with exercise and negatively with blood glucose levels in the BXD mouse genetic reference population (Figure S1B).

**Figure 1.**
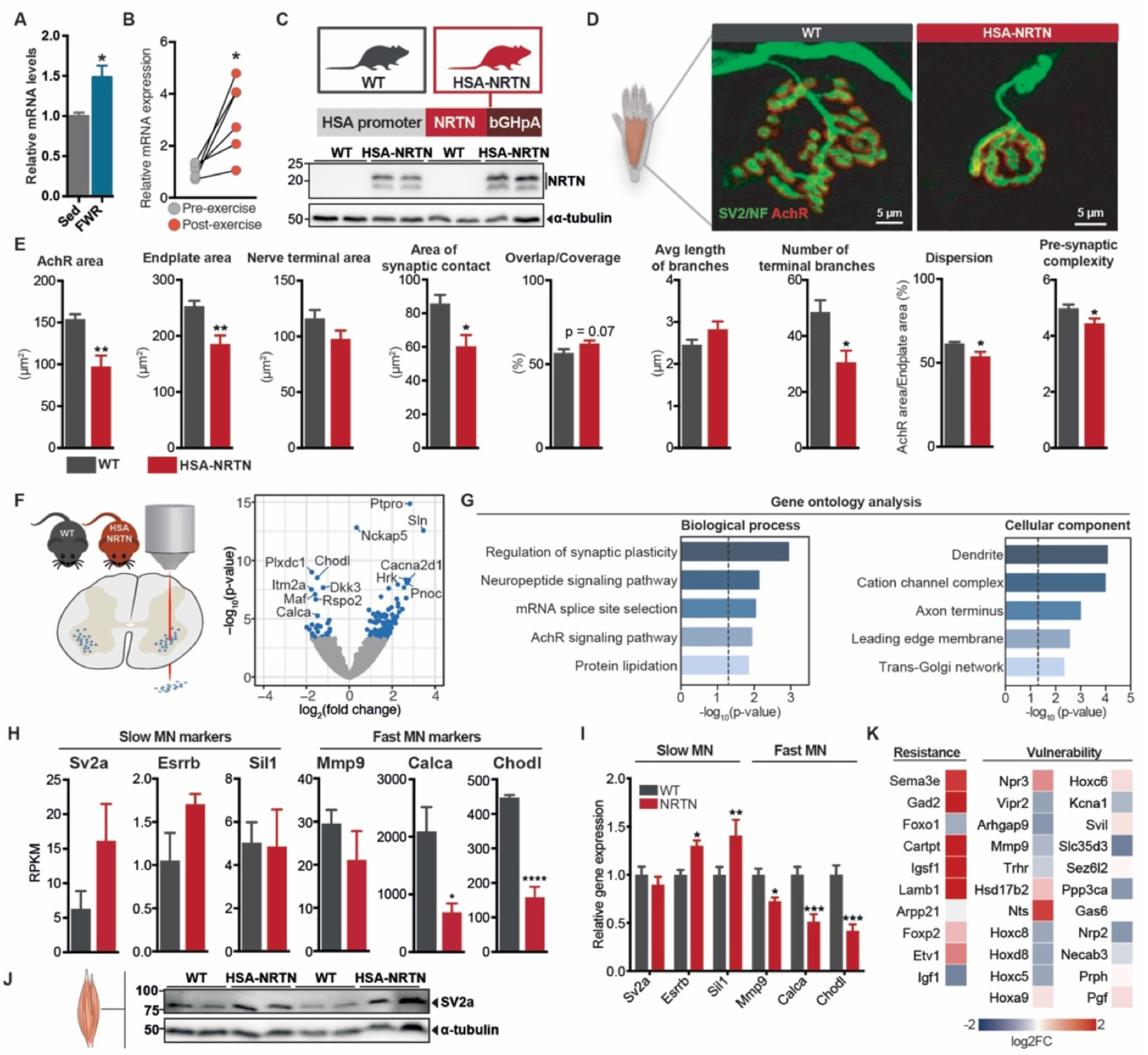
Muscle-derived NRTN promotes motor neuron remodeling. (A) *Nrtn* gene expression in mouse gastrocnemius following 8 weeks of voluntary wheel running (n = 5). (B) *NRTN* gene expression in vastus lateralis muscle of human volunteers after a single session of high-intensity sprint cycling (n=6). (C) Schematic representation of the *Nrtn* expression cassette and NRTN protein levels in gastrocnemius of HSA-NRTN mice and wildtype (WT) littermates. (D) Representative microscopy images of mouse NMJs obtained from FDB whole-muscle mounts of WT and HSA-NRTN mice. MNs were visualized by immunohistochemistry with pan-SV2 and neurofilament (NF) antibodies (shown in green). Acetylcholine receptors (AChRs) were visualized using alpha-bungarotoxin and are shown in red. (E) Quantification of pre-and post-synaptic NMJ properties in WT and HSA-NRTN mice (n = 6). For a detailed flowchart of the NMJ morph analysis platform see Jones et al., 2016. (F) Volcano plot of differential gene expression in lumbar spinal MNs from HSA-NRTN mice following laser-capture microdissection, compared to those of WT littermates (n = 5). (G) Gene ontology analysis of differential gene expression in MNs. (H) Expression of slow and fast MN markers in MNs (n = 5). (I) Gene expression of slow and fast MN markers in the spinal cord of WT and HSA-NRTN mice (n = 5-10). (J) Protein levels of SV2a in gastrocnemius muscle of WT and HSA-NRTN mice. (K) Expression of genes associated with MN susceptibility and resistance to degeneration, in MNs from WT and HSA-NRTN. Bars depict mean values and error bars represent SEM. * p<0.05, ** p<0.01, *** p<0.001, **** p<0.0001. **See also Figure S1**.

To assess the effects of chronically elevated NRTN levels in skeletal muscle, we generated a skeletal muscle-specific *Nrtn* transgenic mouse model (HSA-NRTN). This mouse expresses *Nrtn* under the control of the human alpha-skeletal actin (HSA) promoter (Brennan and Hardeman, 1993), which results in robust *Nrtn* expression in skeletal, and to a lesser extent cardiac, muscles (Figure 1C and S1C). We could not detect NRTN in plasma with the currently available reagents. Contrary to PGC-1α1 (Handschin et al., 2007), transgenic *Nrtn* expression in skeletal muscle did not activate a post-synaptic NMJ gene program (Figure S1D). To evaluate if muscle-derived NRTN affects NMJ morphology, we performed immunohistochemistry analysis of NMJs in whole-muscle mounts of flexor digitorum brevis (FDB) muscle from HSA-NRTN mice and wild-type littermates. Acetylcholine receptor (AChR) aggregates in the post-synaptic membrane were labelled with alpha-bungarotoxin, a potent AchR ligand. MNs were labelled with a pan-synaptic vesicle protein 2 (SV2) antibody combined with a neurofilament (NF) antibody (Figure 1D). Morphometric NMJ analysis revealed that HSA-NRTN mice have overall smaller NMJs, with a marked decrease in AchR cluster area and total post-synaptic endplate area and perimeter but the same number of AChR clusters (Figure 1E and S1E). At the pre-synaptic level, HSA-NRTN had reduced nerve terminal perimeter. The reduced size of both endplates and nerve terminals translated into a significant decrease in the area of synaptic contact, i.e., the total area of overlap between nerve terminals and post-synaptic endplate. The percentage of overlap, defined as the percentage of AChR cluster area in contact with nerve terminals, did not differ significantly from that observed in wild-type mice. Regarding the branching pattern of nerves, HSA-NRTN displayed fewer terminal branches and reduced pre-synaptic complexity, which factors in the number, bifurcations, and length of terminal branches. In sum, the morphometric analysis of HSA-NRTN NMJs suggest a shift towards characteristics of slow-type motor units.

Considering the changes in NMJ morphology observed in the HSA-NRTN mice, we next evaluated to what extent muscle-derived neurturin could elicit retrograde transcriptional remodeling in MNs. To that end, we used laser capture microdissection to capture MN somas from lumbar spinal cords of HSA-NRTN mice and wild-type littermates (Figure S1F-G). Curiously, genotype-blind capture yielded MN pools with different size distributions, with a shift towards reduced soma areas perceivable in HSA-NRTN mice (Figure S1H). We then analyzed the transcriptome of isolated MNs using Smart-seq2 (Figure 1F) (Nichterwitz et al., 2016). The captured cells expressed very high levels of MN markers and low levels of glial markers, confirming their MN identity and negligible glial contamination (Figure S1I). By comparing the transcriptomes of HSA-NRTN and wild-type MNs, we identified a total of 232 differentially regulated genes (Figure 1F and S1J). Among these, the levels of 156 transcripts were higher and 76 were lower in HSA-NRTN MNs, compared to wild-type controls. Gene ontology analysis of differentially expressed genes revealed significant enrichment of genes associated with synaptic plasticity, neuropeptide signaling, and AChR signaling (Figure 1G). Accordingly, this gene set was enriched for transcripts encoding proteins localized to the dendrites, axon terminus and leading edge. Interestingly, among the top genes with reduced expression in MNs from HSA-NRTN mice, we found *Chodl* (Chondrolectin) and *Calca* (Calcitonin-related polypeptide alpha), two known markers for fast MNs (Enjin et al., 2010) (Figure 1F). Metalloproteinase 9 (*Mmp9*), another known marker of fast MN identity (Kaplan et al., 2014), displayed a similar trend. On the other hand, changes in transcript levels of slow MN markers such as synaptic vesicle protein 2a *(Sv2a)* (Chakkalakal et al., 2010) and estrogen-related receptor beta *(Esrrb)* (Enjin et al., 2010), were not statistically significant although trending towards an increase (Figure 1H). We then asked whether the changes in gene expression of MN identity markers were robust enough to be detected by targeted qRT-PCR using bulk lumbar spinal cord RNA. In line with the MN RNA-sequencing data, we detected lower expression of fast MN markers *(Chodl, Calca* and *Mmp9)* and higher expression of slow MN markers *(Esrrb* and *Sil1)* in the spinal cords of HSA-NRTN mice (Figure 1I). Perhaps due to its high expression in other neuronal populations, we did not observe any change in *Sv2a* expression in the spinal cord of HSA-NRTN mice using this approach. Since SV2a is localized to the synaptic vesicles and almost exclusively expressed in neurons, we analyzed SV2a protein levels in gastrocnemius muscle from HSA-NRTN mice and wild-type littermates. HSA-NRTN mice had a two-fold increase in muscle SV2a protein levels compared to wild-type littermates (Figure 1J and S1K). In addition to having different firing properties, fast MNs are also known to be more prone to degeneration under various conditions, such as aging and neuromuscular diseases (Nijssen et al., 2017). Interestingly, HSA-NRTN MNs also expressed higher levels of genes associated with MN resistance to degeneration (Hedlund et al., 2010; Kaplan et al., 2014) (Figure 1K). Conversely, we observed an overall decrease in transcript levels of genes associated with MN susceptibility to degeneration (Figure 1K). Altogether, these data indicate that muscle-derived NRTN promotes a shift towards slow MN identity by eliciting profound transcriptional and morphological changes in MNs and NMJs.

### NRTN induces a shift towards type IIx skeletal muscle fibers

Upon characterization of the HSA-NRTN mice it was evident that their skeletal muscles had a strong red/brown coloration, usually indicative of slow, oxidative fibers (Figure 2A). Skeletal muscles from HSA-NRTN mice were also smaller, with decreased soleus, gastrocnemius and tibialis anterior (TA) mass (Figure 2B). Accordingly, muscle fibers in HSA-NRTN TA showed a 19.1% reduction on average cross-sectional area (CSA), with a higher frequency of CSAs under 2000 μm^2^ and a lower frequency of CSAs above 2500 μm^2^ (Figure 2C-E). The changes in coloration and fiber CSA distribution prompted us to assess whether NRTN triggers a change in skeletal muscle fiber type composition. In line with the shift observed in fiber CSA distribution, HSA-NRTN TA had a substantially higher number of rather compact type IIx fibers, at the expense of bulkier type IIb fibers (Figure 2F-G and S2A). By analyzing the relative abundance of MyHC-IIb and MyHC-IIx proteins by immunoblotting, we observed a similar fiber type switch in gastrocnemius (Figure 2H), which was further substantiated by concomitant changes in the genes encoding MyHC-IIx *(Myh1)* and MyHC-IIb *(Myh4)* (Figure S2B).

**Figure 2.**
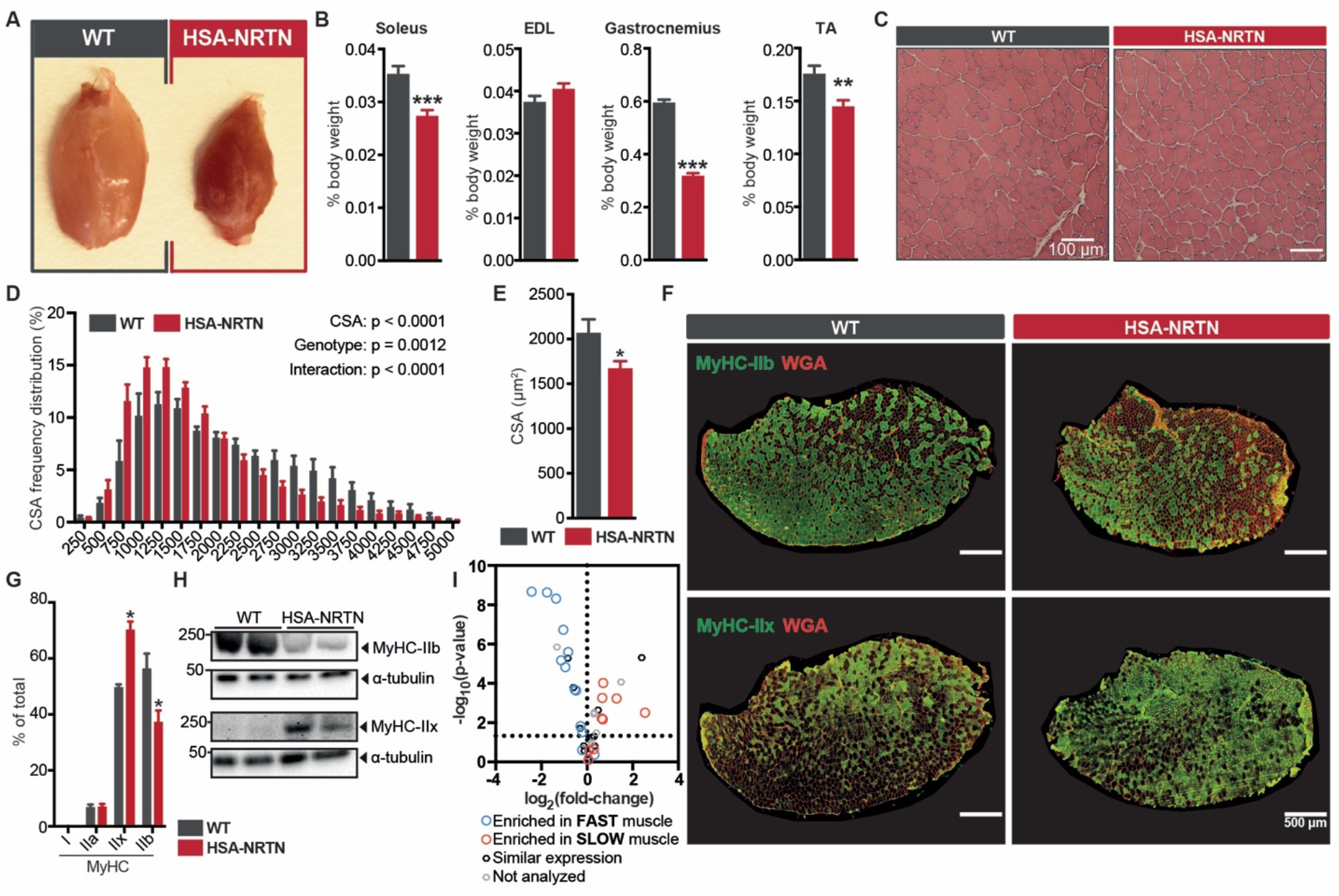
NRTN triggers a fiber type switch in skeletal muscle. (A) Morphology of the gastrocnemius of wild-type (WT) and HSA-NRTN mice. (B) Weight of soleus, extensor digitorum longus (EDL), gastrocnemius, and tibialis anterior (TA) muscles (n = 9-12) normalized by body weight. (C) Histology of WT and HSA-NRTN TA muscles as determined by hematoxylin and eosin staining. (D and E) Fiber cross-sectional area (CSA) distribution (D) and average (E) in TA muscles of WT and HSA-NRTN mice (n = 5-7). (F and G) Immunohistochemical analysis of fiber type composition in TA muscles of WT and HSA-NRTN mice. Cell membranes are visualized using wheat germ agglutinin (WGA, in red). Representative microscopy images (F) and the corresponding quantification (G) are shown. (H) MyHC-IIx and MyHC-IIb protein levels in gastrocnemius muscles of WT and HSA-NRTN mice. (I) Gene expression of calcium handling machinery components. The graph shows the relative expression of genes involved in calcium handling in gastrocnemius muscles of wild-type and HSA-NRTN mice. Each gene is annotated as enriched in fast muscle (EDL) or slow muscle (soleus) based on published gene array data comparing mouse EDL and soleus muscles (Drexler et al., 2012) (n = 6). Bars depict mean values and error bars represent SEM. * p<0.05, ** p<0.01, *** p<0.001. **See also Figure S2**.

In addition to the muscle contractile proteins and MN identity, the third pillar needed for a functional switch in muscle characteristics is the calcium handling machinery (Bassel-Duby and Olson, 2006). Indeed, we observed marked changes in the expression of multiple components of the calcium handling machinery (Figure S2C). When overlaid with published gene array data from the predominantly slow-twitch soleus and the predominantly fast-twitch extensor digitorum longus (EDL) muscles (Drexler et al., 2012), we could determine that NRTN induces the expression of calcium handling components enriched in slow-twitch muscle and represses the expression of those enriched in fast-twitch muscle (Figure 2I). These results show that ectopic NRTN expression in skeletal muscle promotes a shift towards more oxidative and fatigueresistant type IIx fibers.

### NRTN improves mitochondrial function and promotes angiogenesis

To evaluate potential changes in mitochondrial function in skeletal muscle of HSA-NRTN mice, we analyzed NADH dehydrogenase and succinate dehydrogenase (SDH) activities using tetrazolium blue stains. In wild-type muscle, we observed the expected activity pattern for both dehydrogenases, consisting of an intercalated mixture of highly oxidative fibers (deep blue) and fibers with low oxidative capacity (white to light blue) (Figure 3A). Remarkably, almost all the fibers from HSA-NRTN muscle displayed high succinate and NADH dehydrogenase activities (Figure 3A and S3A). Importantly, after segregating fibers with high and low NADH dehydrogenase activity, fibers with similar activities showed no difference in fiber CSA between HSA-NRTN and wild-type mice (Figure S3B). Moreover, the expression of multiple genes commonly associated with skeletal muscle atrophy (atrogenes) were either unchanged or reduced in HSA-NRTN skeletal muscle (Figure S3C-D). Thus, these data indicate that the reduction in fiber CSA in HSA-NRTN mice is due to a shift in the contractile and metabolic properties of the fibers and not to bona fide muscular atrophy. In line with these observations, skeletal muscles from HSA-NRTN mice have higher protein levels of components of all the mitochondrial electron transport chain complexes (Figure 3B and S3E). To evaluate whether supplying exogenous NRTN could improve mitochondrial function in skeletal muscle cells, we treated fully differentiated mouse primary myotubes with recombinant NRTN (rNRTN) and measured mitochondrial respiration. Indeed, myotubes treated with rNRTN displayed increased basal respiration and maximal respiratory capacity (Figure 3C and S3F).

**Figure 3.**
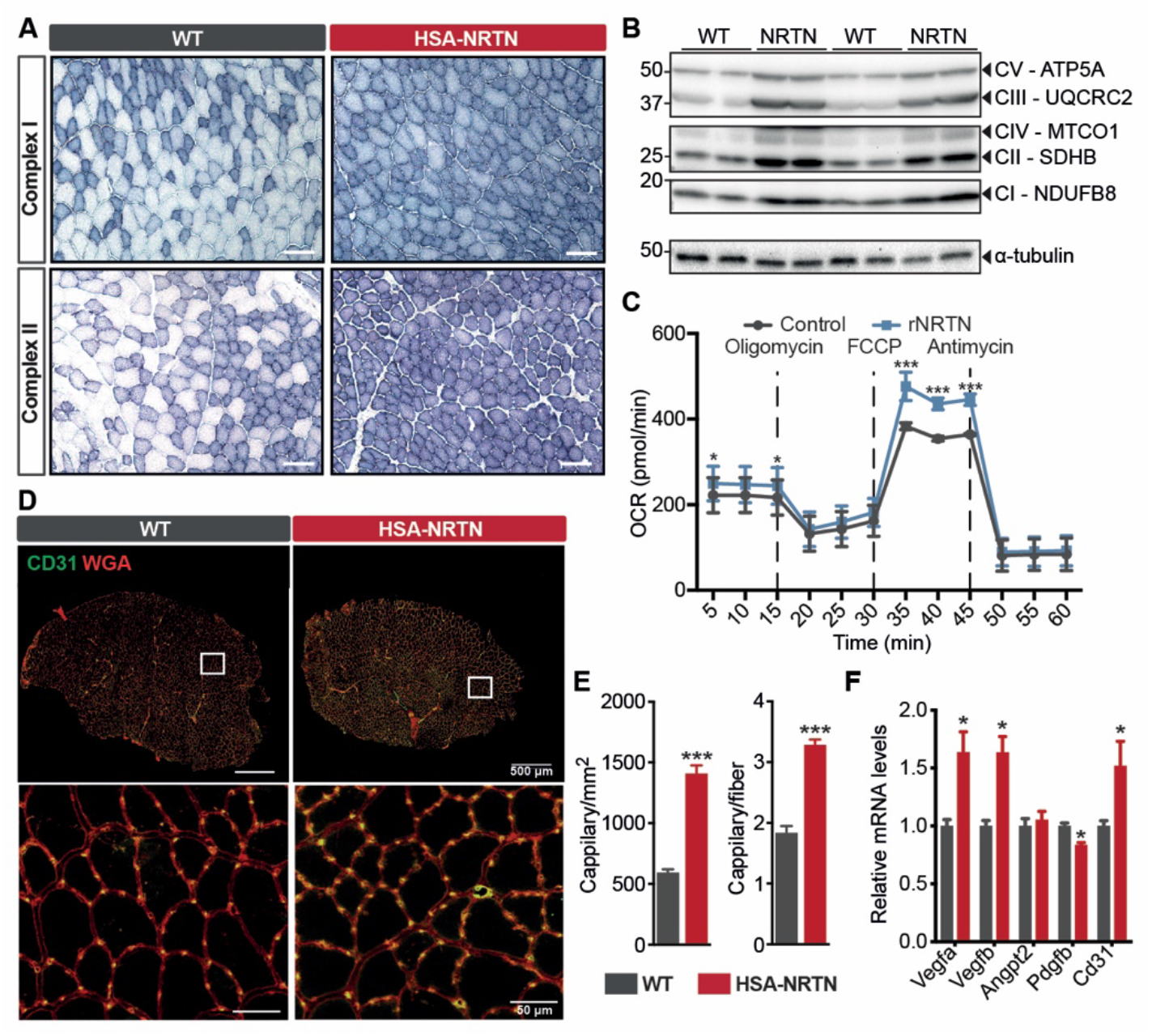
NRTN enhances mitochondrial function and angiogenesis. (A) NADH dehydrogenase (complex I) and succinate dehydrogenase (complex II) activity stains in TA muscles of wild-type (WT) and HSA-NRTN mice. (B) Immunoblots of components of the mitochondrial electron transport chain in gastrocnemius muscles of WT and HSA-NRTN mice. (C) Oxygen consumption rate (OCR) in mouse primary myotubes treated with 10 ng/ml rNRTN (blue) vs vehicle (control, black) for 24h. Baseline OCR and sequential supplementation with 1 μM oligomycin, 1 μM FCCP, and 2 μM antimycin (n = 3). (D) Immunostaining for blood vessels using CD31 in TA muscles of WT and HSA-NRTN mice. (E) Quantification of blood vessel density in TA muscles of WT and HSA-NRTN mice (n = 5-7). (F) Gene expression analysis of angiogenic factors in gastrocnemius muscles of WT and HSA-NRTN mice (n = 6). Bars depict mean values and error bars represent SEM. * p<0.05, ** p<0.01, *** p<0.001. **See also Figure S3**.

Heightened mitochondrial oxidative capacity increases the demand for oxygen and nutrient supply to skeletal muscle. For that reason, we sought to determine if muscle Nrtn expression also activates angiogenesis to increase blood supply to the tissue. By staining skeletal muscle cross-sections from HSA-NRTN mice with an antibody against CD31 we could determine a robust increase in capillary density (Figure 3D), with circa three times more capillaries per square millimeter than wild-type littermates (Figure 3E). To account for the reduced fiber CSA in HSA-NRTN mice, we also normalized the capillary count to the total number of fibers. This revealed a 63% increase in capillaries per fiber in the skeletal muscle of HSA-NRTN mice (Figure 3E). Analysis of HSA-NRTN skeletal muscle gene expression revealed higher levels of the angiogenic factors vascular endothelial growth factor (Vegf) A and B (Figure 3F). These data indicate that NRTN enhances skeletal muscle mitochondrial function and vascularization, thus coupling increased oxidative capacity and oxygen/nutrient supply.

### NRTN promotes opposite transcriptional reprogramming of glucose and fat metabolism

To better understand how NRTN affects skeletal muscle function and metabolism, we performed a global analysis of gene expression by RNA-sequencing of gastrocnemius muscle from HSA-NRTN mice and wild-type littermates. We observed robust transcriptional reprogramming in HSA-NRTN skeletal muscle, with 3418 transcripts induced and 3366 transcripts repressed significantly in relation to wild-type skeletal muscle (Figure 4A and S4A). Gene ontology analysis of differentially expressed genes revealed a strong enrichment for genes involved in energy metabolism, muscle homeostasis/contraction, and inflammatory response (Figure 4B). Consistent with the changes promoted by NRTN on fiber type composition and calcium handling, we observed a strong repression of isoforms of contractile proteins enriched in fast muscles (e.g. myosin binding protein C2, *Mybpc2*), accompanied by an upregulation of isoforms enriched in slow or cardiac muscle (e.g. *Mybpc1*) (Figure S4B). Moreover, by dissecting the pathways involved in energy metabolism, we could determine that transgenic *Nrtn* expression has opposing transcriptional effects on glucose and lipid metabolism (Figure 4C). The expression of the vast majority of glycolytic and glycogenolytic genes was reduced in HSA-NRTN skeletal muscle. Conversely, genes involved in fatty acid uptake (e.g. *Lpl, Cd36*), intracellular transport (e.g. *Fabp3, Cpt1a*) and beta-oxidation (e.g. *Acadm, Acadl*) were induced in HSA-NRTN skeletal muscle, as were most genes encoding enzymes of the TCA cycle. We observed a similar trend for genes encoding the subunits of the various ETC complexes, although with a lower magnitude of fold change and statistical significance (Figure S4C). The transcriptional changes elicited by NRTN on carbohydrate and lipid metabolism are summarized in Figure 4D.

**Figure 4.**
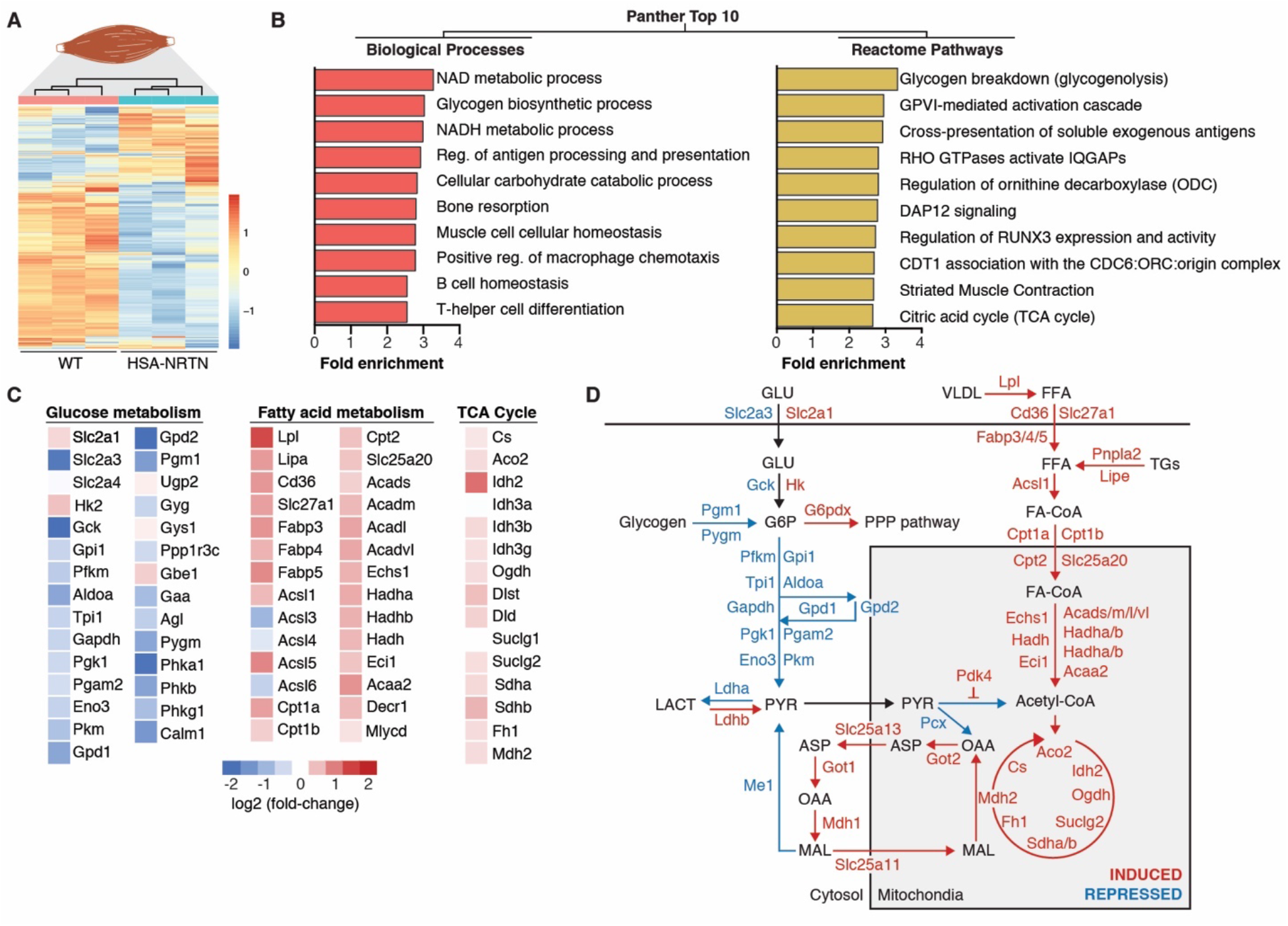
Analysis of HSA-NRTN skeletal muscle by RNA-sequencing. (A) Heatmap summary of changes in gene expression in gastrocnemius muscle of HSA-NRTN mice, compared to wild-type (WT) (n = 3). (B) Gene ontology analysis of genes differentially expressed in HSA-NRTN muscle. (C) Heatmaps showing the relative changes in expression of genes involved in glucose and fatty acid metabolism and the TCA cycle (n = 3). (D) Diagram illustrating the coordinated regulation of genes involved in glucose and fatty acid metabolism in HSA-NRTN skeletal muscle, compared to WT. **See also Figure S4**.

Our results show that the HSA-NRTN mice recapitulate to a large extent the phenotype observed in skeletal muscle-specific PGC-1α1 transgenic models (Arnold et al., 2014; Gill et al., 2018; Handschin et al., 2007; Lin et al., 2002). To ascertain how the transcriptional changes elicited by NRTN in skeletal muscle overlap with those brought about by PGC-1α1, we compared our RNA-sequencing data to published skeletal muscle gene array data from MCK-PGC-1α1 mice (Pérez-Schindler et al., 2012). Strikingly, more than half the genes differentially expressed in MCK-PGC-1α1 (compared to wild-type muscle), were also differentially regulated in HSA-NRTN muscle (Figure S4D). Gene ontology analysis revealed that the overlapping gene panel is enriched for genes involved in various metabolic pathways, including fatty acid oxidation, TCA cycle and glycogen biosynthesis. Pathways involved in excitation-contraction coupling, such as muscle contraction, electrical coupling and calcium release from the sarcoplasmic reticulum, were also overrepresented. The panel of genes differentially expressed exclusively in MCK-PGC-1α1 mice was strongly enriched for mitochondrial processes, suggesting that PGC-1α1 is a stronger activator of mitochondrial biogenesis than NRTN. Despite the strong overlap with the gene programs regulated by PGC-1α1, there were 4472 genes differentially expressed exclusively in HSA-NRTN mice. It is important to consider that the higher sensitivity and dynamic range of RNA-sequencing (compared to gene array used in MCK-PGC-1α1 muscle) may contribute to the high number of differentially expressed genes observed exclusively in the HSA-NRTN gene set. These genes were mainly enriched for processes related to immune cell function, particularly T helper cell lineage commitment. Interestingly, we observed increased expression of markers of anti-inflammatory Th2 and Treg lineages, accompanied by decreased levels of pro-inflammatory Th1 and Th17 markers (Figure S4E). Collectively, these data show that HSA-NRTN muscle displays a transcriptional signature that partly overlaps with that of MCK-PGC-1α1 muscle and that better equips muscle cells to perform fatty acid oxidation and mitochondrial respiration.

### Exercise performance and motor coordination are enhanced in HSA-NRTN mice

Our data indicate that NRTN promotes coordinated changes in MNs and muscle fibers. To assess the implications of these changes on muscle function, we subjected HSA-NRTN mice and wild-type littermates to a series of functional tests. Strikingly, HSA-NRTN showed a remarkable improvement in motor coordination. Not only did HSA-NRTN mice greatly outperform wild-type littermates on the first rotarod trial, they also showed a greater improvement in subsequent trials (Figure 5A). The improvement in rotarod performance was independent of body weight (Figure S5A). In addition, HSA-NRTN were able to run twice as much as wild-type littermates when subjected to a treadmill running capacity test (Figure 5B). On the other hand, HSA-NRTN mice had decreased grip strength in comparison to wild-type littermates, which is consistent with the changes observed in muscle size and fiber type composition (Figure 5C). To further evaluate the effects of NRTN on muscle contractile properties, we analyzed ex vivo contractile force and fatigability in mostly-oxidative soleus and mostly-glycolytic EDL muscles. In HSA-NRTN soleus, we observed a substantial decrease in absolute force production for frequencies over 50 Hz (Figure 5D). This decrease was entirely due to the reduced soleus mass (Figure S5B). In fact, when normalized for CSA (specific force), the soleus from HSA-NRTN mice displayed increased contractile force at 15-30 Hz and reached a plateau similar to that of wild-type mice. On the EDL, HSA-NRTN muscle showed a similar reduction in both absolute and specific force at frequencies over 50 Hz, indicating that the reduction in EDL contractile force was independent of CSA (Figure 5E and S5B). As for fatigability, HSA-NRTN soleus fatigued at the same rate as wild-type soleus but recovered much faster (Figure 5D). On the other hand, the EDL from HSA-NRTN mice retained contractile force for longer and recovered at the same rate as wildtype EDL (Figure 5E). Taken together, these data indicate that muscle-derived NRTN triggers muscle-specific changes on contractile properties which contribute to an overall improvement on motor coordination and exercise performance.

**Figure 5.**
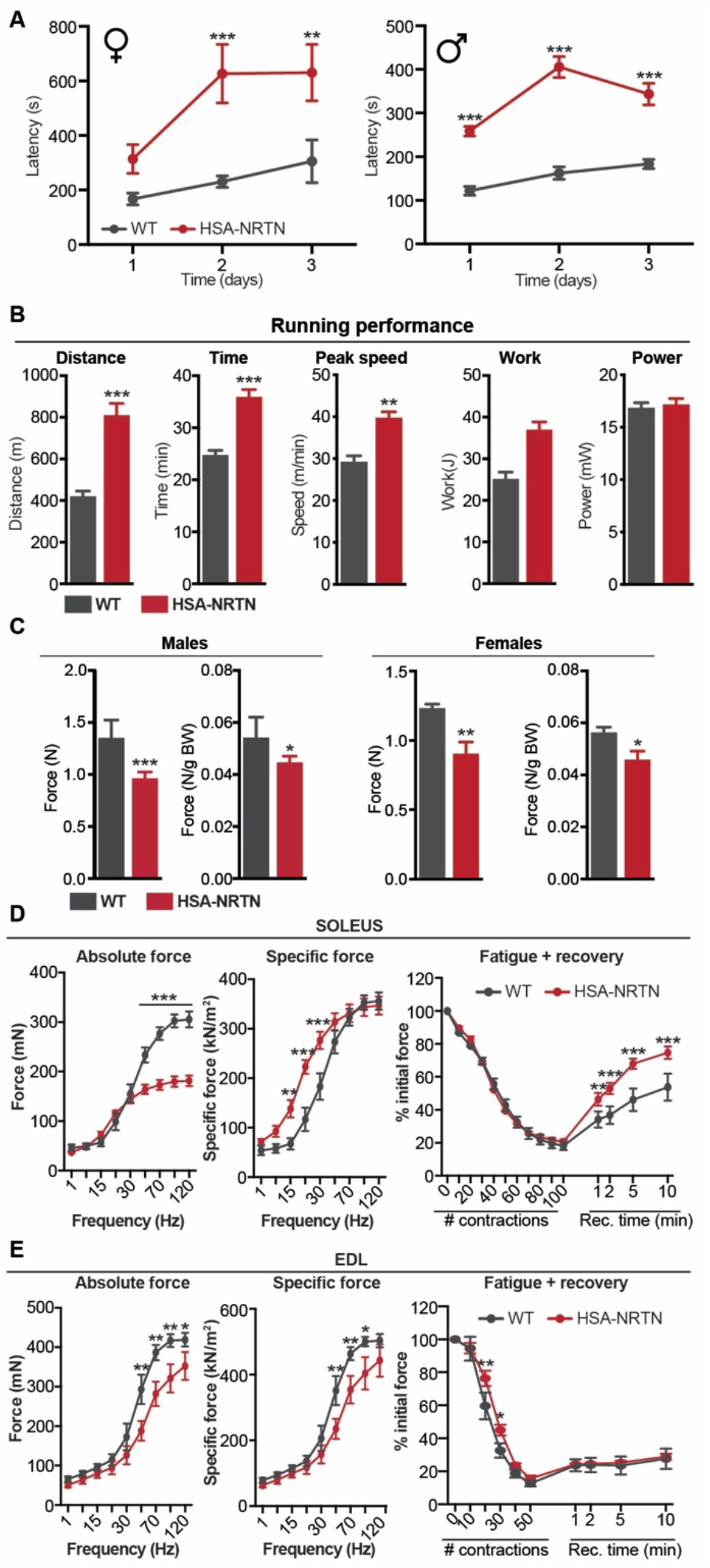
Enhanced motor coordination and running performance in HSA-NRTN mice. (A) Rotarod performance test. Graphs show the maximum time to fall (latency) achieved by wild-type (WT) and HSA-NRTN mice in each of the three rotarod test sessions held in consecutive days (n = 4-11). (B) Total distance and time, peak speed, work executed, and power output achieved by WT and HSA-NRTN mice in a treadmill running performance test (n = 4). (C) Absolute and body weight-normalized grip strength test of WT and HSA-NRTN mice (n = 6-7). (D-E) Ex vivo analysis of contractile force and fatigability of soleus (D) and EDL (E) muscles from WT and HSA-NRTN mice (n = 4). Bars depict mean values and error bars represent SEM. * p<0.05, ** p<0.01, *** p<0.001. **See also Figure S5**.

### Muscle-derived NRTN reduces adiposity and improves glucose homeostasis

The metabolic status of skeletal muscle has extensive repercussions on whole-body energy homeostasis. Given the profound changes triggered by NRTN on skeletal muscle contractile and metabolic properties, we sought to determine the systemic effects of muscle-derived NRTN on energy homeostasis. When compared to wild-type littermate controls, HSA-NRTN mice had lower body weight (Figure 6A) and altered body composition, with a higher percentage of lean mass (Figure 6B) and a lower percentage of fat mass (Figure 6C). We did not observe differences in bone mineral density (Figure 6D) nor in the length of the tibia and femur (Figure S6A). Consistent with the reduction in overall adiposity, HSA-NRTN mice had reduced epididymal white adipose tissue (eWAT) and brown adipose tissue (BAT) mass (Figure 6E-G). To gain further insights into the effects of muscle-secreted NRTN on whole-body energy homeostasis, we monitored HSA-NRTN mice and wild-type littermates in metabolic cages. Food consumption (Figure S6B), locomotor activity (Figure S6C), and respiratory exchange ratio (Figure S6D) were similar for HSA-NRTN and wild-type mice. To account for the pronounced difference in body weight, we performed an analysis of covariance on the various metabolic parameters monitored. By doing so, we did not observe any statistically significant changes in oxygen consumption (Figure 6H), carbon dioxide production (Figure 6I), energy expenditure (Figure 6J), or energy balance (Figure S6E). We also did not observe transcriptional activation of thermogenic gene programs in eWAT (Figure S6F), iWAT (Figure S6G) or BAT (Figure S6H), although the expression of several beige adipocyte markers was elevated in the iWAT of HSA-NRTN mice (Figure S6G). Regarding glucose homeostasis, HSA-NRTN mice displayed improved glucose tolerance (Figure 6K), despite having lower fed plasma insulin levels (Figure 6L) and unchanged fasting plasma insulin levels (Figure S6I). In line with these results, we detected a marked increase in the protein levels of the glucose transporter SLC2A4 (GLUT4) in skeletal muscle from HSA-NRTN mice (Figure 6M). Moreover, we could verify by immunohistochemistry that the increased GLUT4 protein seen in HSA-NRTN muscle is mostly localized at the cellular membrane (Figure S6J). Thus, muscle-derived NRTN promotes metabolic adaptations that result in reduced adiposity and improved glucose homeostasis without an overt adipose tissue browning phenotype.

**Figure 6.**
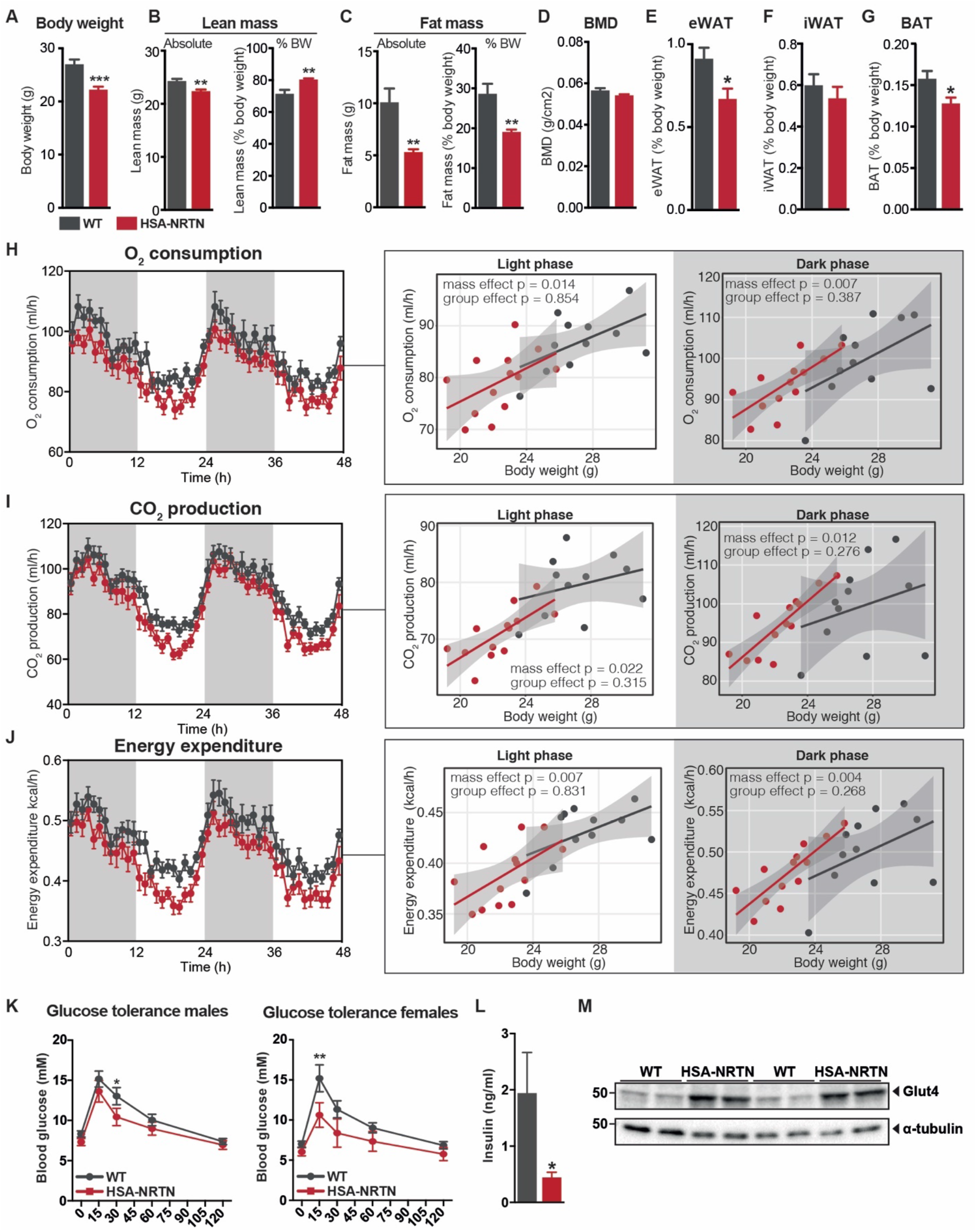
Reduced adiposity and improved glucose homeostasis in HSA-NRTN mice. (A) Body weight of wild-type (WT) and HSA-NRTN mice (n = 11-14). (B-C) Analysis of body composition by Dual-energy X-ray absorptiometry. Graphs display the percentage of lean (B) and fat (C) mass, and bone mass density (BMD, D) of WT and HSA-NRTN mice (n = 8). (E-G) Tissue mass of epididymal white adipose tissue (eWAT, E), inguinal WAT (iWAT, F) and brown adipose tissue (BAT, G) from WT and HSA-NRTN mice, normalized by body weight. (H-J) Metabolic evaluation of WT and HSA-NRTN using a comprehensive laboratory animal monitoring system (CLAMS). Graphs on the left side of each panel represent absolute oxygen consumption (H), carbon dioxide production (I), and energy expenditure (J) in wild-type and HSA-NRTN mice for two days and two nights inside the system. The regression plots on the right side of each panel depict the respective metabolic parameter as a function of mouse weight, in the dark and light cycles (n = 8). (K) Intraperitoneal glucose tolerance test WT and HSA-NRTN mice, following 5 hours of fasting (n = 5-15). (L) Fed-state plasma insulin levels in WT and HSA-NRTN mice (n = 10-13). (M) Immunoblot of Glut4 in gastrocnemius muscles of WT and HSA-NRTN mice. Bars depict mean values and error bars represent SEM. * p<0.05, ** p<0.01, *** p<0.001. **See also Figure S6**.

### NRTN activates RET signaling in skeletal muscle

To investigate the molecular mechanisms that mediate the effects of NRTN on skeletal muscle, we performed activity-based kinase profiling of skeletal muscle from HSA-NRTN mice and wild-type littermates. Compared to wild-type muscle, we observed a significant increase in the phosphorylation of 42 tyrosine kinase substrate peptides and 15 serine/threonine kinase substrate peptides (Figure S7A). Interestingly, one of the top differentially phosphorylated peptides included in the array was a peptide from the RET tyrosine kinase, which functions as a co-receptor for NRTN (Figure S7A). Kinase activity was assessed through computational analysis of the differentially phosphorylated peptide signatures (Figure 7A). Among the kinases more active in HSA-NRTN than wild-type muscle, there were multiple mitogen-activated protein kinases (MAPK), including MAPK1 (ERK2), MAPK3 (ERK1), MAPK8-10 (JNK1-3), and MAPK14 (p38) (Figures 7A and S7B). To better understand how the determined activity of these kinases integrates in a signaling pathway, we constructed a protein-protein interaction network using STRING. As shown in Figure 7B, this yielded a very interconnected network centered around members of the MAPK and SRC families of kinases. Two of the kinases at the core of the network were ERK2 and SRC kinase, both of which are known to signal downstream of the NRTN receptor in neurons (Airaksinen and Saarma, 2002). Interestingly, pathway analysis of differentially active kinases revealed that axon guidance was among the top pathways enriched in the HSA-NRTN muscle kinase network (Figure 7C). Like the main network, the axon guidance sub-network was also centered around SRC kinase and MAPKs, particularly ERK2 and p38 (Figure S7C). In addition to canonical signaling via GFRalpha2:RET and downstream signaling cascades, NRTN can also signal through neural cell adhesion molecule (NCAM) (Paratcha et al., 2003). Notably, kinases downstream of both RET and NCAM displayed higher activity in HSA-NRTN muscle with both branches converging on ERK1/2 (Figure 7D). These data indicate that NRTN can activate the same signaling cascades in skeletal muscle as it does in neurons.

**Figure 7.**
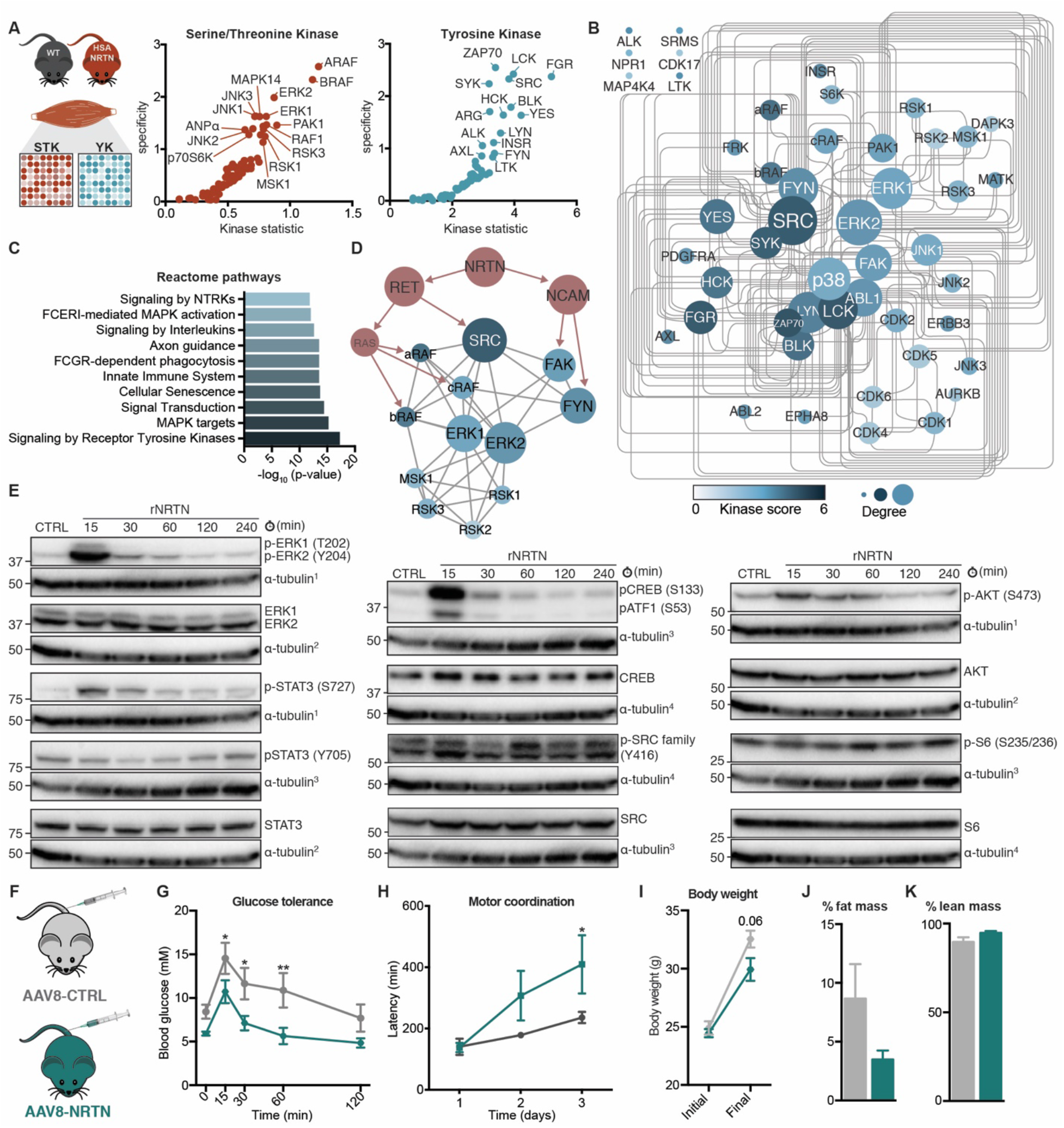
NRTN activates RET signaling in skeletal muscle. (A) Kinase activity profiling by phosphopeptide microarray in wild-type (WT) and HSA-NRTN gastrocnemius muscle extracts. Kinase activity was predicted by computational analysis of differentially phosphorylated peptide signatures. The kinase statistic parameter reflects the change in kinase activity, whereas the specificity parameter indicates how specific the peptide signature is to a particular kinase (n = 3-5). (B) Protein-protein interaction (PPI) network of differentially active kinases. Kinase score reflects kinase activation in HSA-NRTN muscle, compared to wild-type, and the node degree is a measure of how connected each node is within the network. (C) Pathway analysis of differentially active kinases. (D) PPI subnetwork illustrating how the differentially active kinase PPI overlaps with canonical NRTN signaling through RET and NCAM. (E) Immunoblot analysis of mouse primary myotubes treated with 50 ng/ml rNRTN for the indicated times. Shown are representative immunoblots (n = 3). (F-K) Systemic delivery of NRTN to mice. AAV8 vectors encoding a construct expressing *Nrtn* (AAV8-NRTN) or a control construct (AAV8-CTRL) were injected intravenously into C57BL/6N mice. Mice were subjected to glucose tolerance (G) and rotarod (H) tests 12 weeks after. Body weight was recorded (I) and the percentage of fat (J) and lean (K) mass was analyzed by magnetic resonance imaging (n = 5). Bars depict mean values and error bars represent SEM. * p<0.05, ** p<0.01. **See also Figure S7**.

Our results suggest that NRTN signals to both MNs and muscle fibers. Indeed, *Gfra2* (NRTN’s main receptor)(Airaksinen and Saarma, 2002) was expressed in cultured mouse primary myotubes and its expression increased throughout myotube differentiation (Figure S7D). *Gfra1,* the main receptor for GDNF, displayed the opposite expression profile, whereas RET expression was stable during myotube differentiation. To test whether NRTN can act directly on muscle cells, we treated fully differentiated mouse primary myotubes with rNRTN and analyzed protein phosphorylation by immunoblotting at various times points. Treatment with rNRTN induced robust, fast, and transient phosphorylation of ERK1/2 and their downstream target cAMP-responsive element-binding protein (CREB) but had modest effects on the phosphorylation status of SRC family kinases (Figure 7E). In addition to the RAF/MEK/ERK pathway, RET activates multiple other downstream effectors, including AKT and signal transducer and activator of transcription 3 (STAT3). In cultured myotubes, treatment with rNRTN activated AKT and induced STAT3 phosphorylation specifically at serine 727, which is mostly associated with its mitochondrial activity (Luo et al., 2016b; Yang and Rincon, 2016). Interestingly, rNRTN simultaneously reduced STAT3 tyrosine phosphorylation, which is mainly associated with its nuclear transcriptional activity. These results indicate that, in addition to neurons, NRTN can also activate signaling downstream of its receptors in skeletal muscle cells.

Muscle-derived NRTN showed clear benefits on muscle physiology and systemic metabolism, which can be of therapeutic value. To evaluate whether similar benefits can be triggered by systemic NRTN delivery, we performed intravenous injections of adeno-associated viral vectors (serotype 8, AAV8) to deliver a fulllength *Nrtn* construct (or control construct) to the liver (Figure 7F). This method achieves robust expression in liver and secretion to plasma (Rao et al., 2014; Sands, 2011). Mice injected with AAV8-NRTN had improved glucose tolerance (Figure 7G), compared to mice injected with control AAV8. Strikingly, AAV8-NRTN injection also improved rotarod performance (Figure 7H). We did not observe significant differences in body weight (Figure 7I) or composition (Figure 7J-K), although mice injected with AAV8-NRTN tended to have reduced body weight, adiposity, and white adipose tissue mass (Figure S7E). Although we could not consistently detect NRTN in circulation with commercially available reagents, we detected increased levels of mature NRTN in skeletal muscle of mice injected with AAV8-NRTN, despite a reduction in skeletal muscle *Nrtn* expression (Figure S7F-G). In addition, we observed an increase in the phosphorylation of ERK1/2 (Figure S7F), which we show to be robustly activated by NRTN in muscle cells (Figure 7E). These data indicate that NRTN, delivered systemically, can target skeletal muscle and improve motor coordination and glucose homeostasis.

## DISCUSSION

Activating PGC-1α1 in skeletal muscle could open novel therapeutic opportunities for a myriad of muscular, metabolic, and mood disorders. For this reason, many efforts have been made to find ways to activate PGC-1α1 or mimic its biological effects (Arany et al., 2008b; Pettersson-Klein et al., 2018; Zhang et al., 2013), which have been hindered by the fact that transcriptional coactivators are notoriously challenging to target therapeutically. In addition, systemic administration of a PGC-1α1 activator could induce hepatic gluconeogenesis (Sharabi et al., 2017) or cause cancer-related concerns (Luo et al., 2016a). Therefore, myokines downstream of PGC-1α coactivators have recently emerged as appealing alternatives to bypass these hurdles and activate specific aspects of PGC-1α-related biology. To date, most studies have identified PGC-1α-driven myokines with endocrine effects, particularly those that improve systemic metabolism by promoting browning of adipose tissue (Boström et al., 2012; Rao et al., 2014; Roberts et al., 2014). Some of the few known PGC-1α-related myokines with direct effects on the muscle fiber (myostatin and IGF1) are controlled by PGC-1α4, a PGC-1α variant linked to muscle hypertrophy (Ruas et al., 2012). The effects of NRTN on systemic metabolism seem to stem mostly from local effects on skeletal muscle, achieved through autocrine and paracrine signaling to myofibers and MNs, respectively. In fact, NRTN interacts with heparan sulfates in the extracellular matrix (Runeberg-Roos et al., 2016; Sandmark et al., 2018), which limits its ability to reach circulation upon secretion from muscle. This is a common characteristic of GDNF family ligands and many growth factors and is often important for biological function, by increasing local ligand concentrations and facilitating ligand-receptor interactions (Bespalov et al., 2011). Although many heparan sulfate-binding factors, including GDNF, are detectable in circulation (Roh and So, 2017), we were unable to detect NRTN in plasma using commercially available reagents. In addition, NRTN does not increase energy expenditure nor does it activate a thermogenic gene program in adipose tissue, further indicating that NRTN acts through mechanisms that diverge from other known PGC-1α-driven myokines.

Our data demonstrate that myofiber-derived NRTN promotes a shift in MN specification, towards a slow MN identity. MNs from HSA-NRTN mice display smaller cell bodies and reduced nerve terminal branching, and formed smaller, less complex NMJs, all of which are common morphological features of slow MNs (Kanning et al., 2010). These features partially overlap with those observed in MCK-PGC-1α1 mice, which display reduced AChR aggregate area but have higher nerve terminal branching and NMJ complexity (Arnold et al., 2014). In addition, unlike PGC-1α1 (Handschin et al., 2007), NRTN did not activate a myocellular post-synaptic gene program, indicating that its NMJ remodeling effects are mediated mainly by retrograde signaling to MNs. This notion is further substantiated by the changes observed in the transcriptional profile of spinal MNs from HSA-NRTN mice, with increased expression of slow MN markers and decreased expression of fast MN markers. Importantly, locomotion was unchanged in HSA-NRTN mice, which indicates that this shift was not due to increased neuromuscular activity.

The notion that myofibers can exert direct influence on MNs is strengthened by various studies that elucidated the effects of myocellular adaptations and myofiber-derived neurotrophic factors (NFs) on NMJ formation, maintenance, and remodeling (Chakkalakal et al., 2010; Delezie et al., 2019; Nguyen et al., 1998). Importantly, different NFs elicit specific changes on the NMJ and muscle physiology, which indicates that despite having overlapping functions in supporting neuronal survival and tissue innervation, NFs exert non-redundant roles in motor neuron specification and NMJ remodeling. For example, glial cell line-derived neurotrophic factor (GDNF) is elevated by physical exercise in skeletal muscle and correlates positively with NMJ endplate area (Gyorkos and Spitsbergen, 2014). Transgenic expression of *Gdnf* in muscle fibers results in larger motor units and hyperinnervation of NMJs (Nguyen et al., 1998; Zwick et al., 2001). Despite signaling through the same coreceptor (RET) and promoting motor neuron recruitment to myotubes in vitro (Mills et al., 2018), myofiber-derived NRTN did not induce skeletal muscle hyperinnervation. This is in agreement with previous reports, showing that postnatal subcutaneous administration of GDNF (but not NRTN) promotes hyperinnervation of NMJs (Keller-Peck et al., 2001) and that genetic ablation of *Gfra1*(favored receptor for GDNF) but not *Gfra2* (favored receptor for NRTN) jeopardizes MN survival (Garcè et al., 2000). Recently, and in sharp contrast with NRTN, brain-derived neurotrophic factor (BDNF) secreted from muscle fibers was shown to promote glycolytic fiber-type specification in skeletal muscle (Delezie et al., 2019). Skeletal muscle-specific *Bdnf* knockout mice show reduced NMJ endplate volume, a transition to type IIx muscle fibers, and increased running capacity. Thus, myofiber-derived NRTN and BDNF elicit virtually opposite effects on motor unit specification and skeletal muscle metabolic properties. The overall neuromuscular adaptive response to stressors such as exercise or denervation likely reflects the combined action of these and other neurotrophic and growth factors, which can be skewed towards a specific outcome depending on their local relative abundance and receptor availability. The fact that the activation and action of NFs is strongly dependent on spatially delimited cues, may explain why NRTN expression in whole adult skeletal muscle tissue is low. Although we could observe increased NRTN expression with exercise on whole muscle samples, it is possible that NRTN expression is activated in select fibers with active NMJ remodeling.

Slow MNs typically innervate type I muscle fibers. However, and despite an overall transcriptional signature of slow myofibers in terms of calcium handling and structural machinery, the main fiber type switch in HSA-NRTN mice was from IIb to IIx fibers, with expression of Myh7 (which encodes MyHC-I) elevated only in the soleus. Interestingly, a similar conversion of IIb to IIx fibers stands out as a common feature in various genetic mouse models with enhanced muscle oxidative metabolism and endurance exercise capacity, such as skeletal muscle-specific PGC-1α1, PGC-1β, or ERRγ transgenic mice (Arany et al., 2007; Fan et al., 2018; Gill et al., 2018). Type IIx fibers are fast-twitch fibers, with mechanical properties in between type IIa and IIb fibers, but are often highly oxidative. Curiously, a large proportion of highly oxidative type IIx fibers is recurrently observed in sprinting wild animals and may allow animals to sustain high running speeds for long periods of time (Curry et al., 2012). Importantly, in MCK-PGC-1α1 mice, approximately one third of type IIx/IIb fibers are innervated by slow SV2a-positive nerve terminals (a 5-fold increase over wild-type controls) (Chakkalakal et al., 2010). To our knowledge, in almost two decades of research, no abnormalities on locomotion, coordination, or endurance were ever described in these animals, which clearly indicates that this apparent MN/myofiber mismatch is fully functional (Gill et al., 2018; Lin et al., 2002). Indeed, HSA-NRTN mice not only did not display any locomotor abnormalities but greatly outperformed wild-type littermates in both running capacity and motor coordination.

We initially identified NRTN as a myokine involved in retrograde signaling to MNs (Mills et al., 2018), but our results indicate that NRTN can also signal to muscle cells, suggesting that myofiber-derived NRTN has both paracrine and autocrine effects. This is supported by the fact that cultured primary myotubes express NRTN’s receptor *(Gfra2)* and co-receptor (Ret) and are responsive to exogenous NRTN. Indeed, treatment of primary myotubes with rNRTN robustly activated signaling cascades downstream of the Gfra2/Ret and increased mitochondrial respiration. The overall product of NRTN signaling simultaneously to MNs and myofibers was a striking increase in skeletal muscle vascularization and oxidative capacity. This oxidative shift is supported by the coordinated induction of gene programs related to fatty acid transport and oxidation and mitochondrial oxidative metabolism, accompanied by an overall reduction in glycolytic transcripts. However, we did not observe any changes on RER, which indicates that the transcriptional reprogramming triggered by NRTN renders muscles more capable of utilizing fatty acids but does not change resting substrate preference.

Although our initial approach was centered on understanding how PGC-1α1 promotes NMJ remodeling, transgenic expression of *Nrtn* in skeletal muscle was sufficient to recapitulate many of the known effects of skeletal muscle PGC-1α1, including but not limited to NMJ remodeling. Given the metabolic phenotype elicited by NRTN, which renders mice leaner and more glucose tolerant, the most obvious potential application of NRTN is perhaps in the context of metabolic disease. One interesting observation though, is that spinal MNs from HSA-NRTN mice expressed higher levels of transcripts associated with MN resistance to degeneration and lower levels of those associated with MN susceptibility to degeneration (Hedlund et al., 2010; Kaplan et al., 2014). Combined with the improvements on motor coordination seen in HSA-NRTN mice, NRTN might also prove useful in the context of neuromuscular diseases, particularly amyotrophic lateral sclerosis (ALS). Indeed, AAV2-NRTN delivered intraspinally to SOD1G93A mice protects MNs and NMJ maintenance, although this effect is limited to the site of injection (Gross et al., 2020). Further supporting the therapeutic utility of NRTN in the context of metabolic disease, chronic administration of rNRTN was recently shown to increase pancreatic insulin content and beta cell mass, and to prevented deterioration of islet organization in diabetic Zucker rats (Trevaskis et al., 2017). This improved glycemic control and prevented the development of hyperglycemia. Interestingly, NRTN does not seem to act directly on beta cells or pancreatic islets, which led the authors to speculate that NRTN may act by improving innervation on a yet unidentified metabolic tissue. In light of our data, we speculate that skeletal muscle might be that tissue, although the contribution of other metabolically active tissues cannot be excluded.

The strong affinity to heparan sulfates limits NRTN exposure and hinders its direct therapeutic utilization. In fact, limited tissue exposure has been suggested as a major contributor to the poor therapeutic efficacy of NRTN in clinical trials for Parkinson’s disease (Marks et al., 2008, 2010; Runeberg-Roos et al., 2016). To try to bypass the limitations imposed by its limited exposure, we used AAV8-NRTN delivery to the liver to achieve continuous systemic NRTN delivery. By having continuous production and secretion of high levels of NRTN by the liver, we aimed to maximize the amount of NRTN that escapes the heparan sulfate sink and reaches circulation. This strategy has been previously used to study other myokines (Rao et al., 2014). With this approach we were able to show that systemically-delivered NRTN can target skeletal muscle and improve motor coordination and glucose tolerance. The recently resolved crystal structure of NRTN and its receptors, and the subsequent identification of heparan sulfate-binding sites on NRTN, paves the way for the development of engineered NRTN variants with improved pharmacokinetic properties (Bigalke et al., 2019; Sandmark et al., 2018). Indeed, NRTN variants with reduced heparan sulfate-binding show increased exposure in vivo and can still signal through Ret kinase in vitro (Runeberg-Roos et al., 2016; Sandmark et al., 2018). It is yet unknown whether these variants retain full signaling capabilities and biological function in vivo, but they establish a good framework for the development of NRTN-based therapeutic agents.

## EXPERIMENTAL PROCEDURES

### Animal Experiments

Mice were housed in groups (3-5 mice per cage) on a 12h light/dark cycle and with ad-libitum access to water and standard rodent chow diet unless otherwise indicated. All experiments were approved by the regional animal ethics committee of Northern Stockholm, Sweden.

### Generation of a muscle-specific NRTN transgenic mouse model

The coding sequence of mouse Nrtn was cloned in front of a 2.4 kb fragment of the human skeletal alpha-actin (HSA; ACTA1) (Brennan and Hardeman, 1993), followed by a 225 bp fragment of the bovine growth hormone polyA sequence. The expression cassette was purified and delivered via pronuclear injection in mouse zygote with a C57bl/6N genetic background. Transgenic mice (HSA-NRTN) were kept as heterozygotes and housed in standard conditions.

### Grip strength test

Forelimb grip strength was measured using a grip strength meter (Bioseb). Mice were held by the tail and allowed to hold the grid of the apparatus with the front paws. Mice were gently pulled away from the grid until they released. The force exerted was measured. Grip strength was measured in triplicate on three consecutive days. Mice were weighed before the first measurement.

### Glucose tolerance test

Mice were fasted for 5 hours, weighted and injected intraperitoneally with 2 mg/kg glucose. Blood glucose levels were measured before and 15, 30, 60, and 120 minutes after glucose administration using a OneTouch Accu-Check glucometer.

### Treadmill performance test

Mice were acclimated to the treadmill (Columbus Instruments, OH, USA) for four consecutive days prior to the treadmill performance test. Acclimation consisted of 5 min of running at 6 m/min followed by 5 minutes at 6-12 m/min. For performance evaluation, mice warmed up for 3 min at 6 m/min. After this point, speed was increased by 3 m/min every 3 minutes until exhaustion was reached. Mice were encouraged to continue running by gentle prodding and classified as exhausted when they could no longer keep running despite prodding. Treadmill inclination was set at 10 degrees for both acclimation and testing sessions. Mice were weighed before the treadmill performance test was performed.

### Metabolic phenotyping by CLAMS

For metabolic phenotyping, mice were placed in individual metabolic cages in a CLAMS (Comprehensive Laboratory Animal Monitoring System, Columbus Instruments, OH, USA) for five consecutive days, with free access to food and water. Food and water consumption, oxygen consumption, carbon dioxide production, and movement were monitored in real-time. On the last night in the system, mice were fasted for 12h. Access to food was reestablished the following morning. Bodyweight was recorded before and after mice were placed in the CLAMS. Data analysis was performed using CalR (Mina et al., 2018).

### Analysis of body composition by DEXA scan

Mice were placed under isoflurane anaesthesia and body composition was analyzed by dual-energy X-ray absorptiometry (DEXA) using a Lunar PIXImus densitometer (GE Medical Systems). Tibia and femur lengths were measured from DEXA scan images using ImageJ.

### Voluntary wheel running

Wild-type C57bl/6J mice were single-housed in cages with free access to a vertical running wheel for 8 weeks. Sedentary controls were single-housed in similar cages but without access to a running wheel. The wheel was locked for 24h before the mice were euthanized.

### Rotarod

Motor coordination was assessed with a rotarod test, performed on a rotating rod (Ugo Basile) that accelerated from 4 to 40 rpm over the course of 5 minutes (an increase of 1 rpm approximately every 8.3 seconds). This equates to a linear speed ranging from circa 0.6 m/min to a maximum of 6 m/min. The time spent on the rod (before falling) per trial was recorded. Animals were tested for 3 consecutive days with 3 trials per day, separated by intervals of at least 15 minutes. The longest duration per day is reported for each animal. All behavioural studies were carried out with the experimenter blind to genotype.

### Ex vivo analysis of muscle force and fatigability

Skeletal muscle force production and fatigue resistance were measured in soleus and EDL muscles. The muscles were kept in a Tyrode solution (121 mM NaCl, 5 mM KCl, 1.8 mM CaCl2, 0.4 mM NaH2PO4, 0.5 mM MgCl2, 24 mM NaHCO3, 0.1 mM EDTA, and 5.5 mM glucose) and mounted between a force transducer in a stimulation chamber with proximal and distal tendons tied to two hooks. The chamber temperature was kept at 31°C with a circulatory water bath and the pH of Tyrode solution was kept at 7.4 by continuous superfusion with carbogen (95% O2, 5% CO2). Muscles were adjusted to optimal length. Force-frequency relationships were determined by stimulating muscles at different frequencies (1Hz to 120 Hz, 1000 ms tetanic duration for the soleus muscles; 1Hz to 150 Hz, 300 ms tetanic duration for the EDL muscles). To calculate muscle specific force (kN/m2), the muscle CSA was calculated by dividing muscle mass by the product of muscle length and muscle density (1.06 g/cm3). To measure fatigue resistance, tetanic force was measured at 70 Hz stimulation with 600 ms duration and 2 second intervals for 100 contractions in the soleus or at 100Hz with 300 ms duration and 2 second intervals for 50 contractions in the EDL. Then the recovery was measured by allowing the muscles to recover for 10 minutes and tetanic force was measured at 1, 2, 5, and 10 minutes after the fatigue protocol.

### AAV-mediated gene delivery to liver

Ten-week-old male C57BL/6N were received 2×10^11^ viral genomes of AAV8 in PBS (empty or expressing full-length mouse *Nrtn*), injected in the tail vein. Mice were group-housed in standard conditions (12h light/dark cycle; ad-libitum access to water and standard rodent chow diet). Glucose tolerance tests and MRI were performed 10 weeks after AAV administration. Rotarod tests were performed 11 weeks after AAV administration. Mice were terminated 12 weeks after AAV administration.

### Determination of plasma insulin levels

Plasma insulin was measured using the Ultra Sensitive Mouse Insulin ELISA Kit (Crystal Chem, IL, USA) according to the manufacturer’s instructions for the low range assay.

### NMJ immunohistochemistry and morphological analysis

FDB muscles were dissected and fixed in 4% formaldehyde for 10 min. Following three washes with PBS, the muscles were incubated in PBS supplemented with 0.1M glycine and 2% BSA for 30 min, washed with PBS, permeabilized for 30 min (2% Triton X-100 in PBS) and blocked for 1 hour (4% BSA, 0.2% Triton X-100 and 5% normal goat serum in PBS). Muscles were then incubated with primary antibodies raised against neurofilament (2H3) (1:100) and synaptic vesicle glycoprotein (pan-SV2) (1:100) (both from the Developmental Studies Hybridoma Bank, University of Iowa, USA) for 48h at 4°C. After 5 washes, muscles were incubated with α-bungarotoxin (Alexa Fluor 594-conjugated, 1:1000, Thermo Fisher Scientific) for 1h and finally with secondary antibodies in PBS for 2h. Samples were then washed with PBS and wholemounted on slides with Vectashield DAPI mounting medium.

Images were acquired and analyzed according to a standardized workflow previously described by Jones et al., 2016. Briefly, z-stack images of individual en-face NMJs with 1um intervals were collected using a Zeiss LSM 710 confocal microscope. Images were analyzed using ImageJ/Fiji (and freely available plugins) to measure various individual pre- and post-synaptic morphological variables, following the NMJmorph workflow (Jones et al., 2016). For the analysis, ≥ 30 NMJs per muscle were analyzed and the mean values were used to plot the data and for the subsequent statistical analysis.

### Laser capture microdissection and RNA-seq of motor neurons

Lumbar spinal cords were dissected and immediately snap-frozen in 2-methylbutane (Sigma-Aldrich) cooled in liquid nitrogen. Spinal cords were embedded in OCT, cryosectioned at −20°C (12 μm coronal sections) onto PEN membrane glass slides (Zeiss) and subsequently stored at −80°C until further processing. The sections were then used for laser capture microdissection of spinal motor neurons as described in Nichterwitz 2016 (Nichterwitz et al., 2016). Briefly, the sections were thawed 30 seconds, subjected to quick histological staining (Histogene, Arcturus/LifeTechnologies) and cells were captured using the Leica LMD7000 system at 40x magnification. Only cells with a visible nucleus with nucleolus and with an area of more than 200 μm2 were captured. 50 cells per sample were collected into the cap of PCR tubes, mixed with 5 μl lysis buffer (0.2% Triton X-100, with 2 U/μl recombinant RNase inhibitor, Clontech) at the end of the collection and were immediately snap-frozen on dry ice. The duration from thawing the slides until freezing of the lysed cells never exceeded 2 hours. Samples were stored at −80°C until library preparation. The libraries were prepared as described in detail in Nichterwitz 2018 (Nichterwitz et al., 2018). Samples were loaded onto an Illumina HiSeq 2500 High-output v4 flow cell and sequenced in a 1×50 bp single read format. Quality control of raw reads was determined using FastQC tool kit (Babraham Bioinformatics, http://bioinformatics.babraham.ac.uk/projects/fastqc). The reads were then aligned with reference genome of Mus Musculus (GRCm38.p13) downloaded from NCBI using STAR aligner tool (Dobin et al., 2013) and reads aligning to gene exons were counted using Featurecounts program(Liao et al., 2014). List of differentially expressed genes (DEGs) between wild-type and NRNT motorneurons was obtained by analyzing raw counts using DESeq2 package (Love et al., 2014) and RPKM values were calculated using *fpkm* function in DESeq2 package. All genes showing p-value (adjusted by Benjamini-Hochberg method) < 0.1 were used in subsequent analyses. The data files were submitted to GEO database: GSEXXXXXX codes will be available before publication.

### Primary myotube cultures and extracellular flux analysis

Mouse primary myoblasts were isolated, maintained, and differentiated as previously described (Rando and Blau, 1994). Fully differentiated myotubes were treated with 50 ng/ml recombinant mouse NRTN (rNRTN, R&D Systems) for 15, 30, 60, 120, or 240 minutes. Cells were quickly washed with PBS and lysed with SDS lysis buffer. For analysis of mitochondrial respiration, fully differentiated myotubes were treated for 24h with 10 ng/ml rNRTN before being subjected to extracellular flux analysis (XF24, Seahorse Bioscience). The mitochondrial stress test assay was performed with 1 mM pyruvate and 5 mM glucose. Oxygen consumption rate (OCR) was measured every 7 minutes. Following baseline measurements, media was sequentially supplemented with 1 uM oligomycin, 1 uM carbonyl cyanide-4-(trifluoromethoxy) phenylhydrazone (FCCP) and 2 uM antimycin.

### Kinase activity profiling arrays

Kinase activity was analyzed in gastrocnemius whole-cell protein lysates using Pamgene’s tyrosine kinase (PTK) and serine/threonine kinase (STK) PamChip arrays, according to the manufacturer’s instructions. Briefly, muscle whole-cell protein lysates were diluted in MPER buffer (Sigma-Aldrich) supplemented with protease and phosphatase inhibitor cocktails. Each assay was performed in duplicate, using 5 μg of protein lysate. Chips were placed in a PamStation 12 System, blocked with 2% BSA and incubated with sample and detection reagents following Pamgene’s recommended protocols. Quantification of spot images, quality control, statistical analysis and further upstream kinase analysis was performed by Pamgene using BioNavigator and in-house analysis pipelines.

Protein-protein interaction networks were generated using STRING (string-db.org; Szklarczyk et al., 2018) and visualized and edited in Cytoscape. Only upstream kinases with a specificity score higher than 0.5 and a mean significance score higher than 1 were used for network generation (total of 48 kinases). Networks were built with the stringency set to 0.7 and a force-directed layout based on score. The distance of nodes situated on the edges of the network was manually adjusted for space optimization. Pathway analysis was performed using STRING. Tile proteomaps were generated from the same set of 48 putative upstream kinases using Proteomaps (Liebermeister et al., 2014).

### Analysis of gene expression

Total RNA was isolated from cell cultures and frozen tissue using TRI reagent (Sigma-Aldrich) according to the manufacturer’s instructions. One μg of RNA was treated with Amplification Grade DNAse I (Thermo Scientific) and 500 ng of DNAse-treated RNA were used for cDNA preparation using the High-Capacity cDNA Reverse Transcription Kit (Applied Biosystems). Quantitative real-time PCR was performed in ViiA7 and QuantStudio 6 real-time PCR systems (Applied Biosystems) using the SYBR Green PCR Master Mix (Applied Biosystems). Relative gene expression was calculated using the ddCt method and normalized to the expression of TATA-binding protein (Tbp).

### Global gene expression analysis of skeletal muscle by RNA-sequencing

Total RNA was extracted from gastrocnemius muscles from wild-type and HSA-NRTN muscles using TRI reagent (Sigma-Aldrich) according to the manufacturer’s instructions. RNA was further cleaned and treated with DNAse using NucleoSpin RNA II columns (Machery Nagel). RNA integrity was verified using an Agilent Bioanalyzer. RNA-sequencing was performed at GATC Biotech (Konstanz, Germany). Libraries were prepared with the Illumina TruSeq Stranded mRNA Library Preparation Kit using a polyA-enrichment strategy. Quality control of raw reads was performed using the FastQC tool kit (Babraham Bioinformatics). The clean sequenced reads were aligned to the mm10 mouse genome using HISAT2 (Kim et al., 2015) and feature count was performed using HTSeq framework (Anders et al., 2015). Differential expression between wild-type and HSA-NRTN muscle was determined using the DESeq2 package by applying regularized log transformation (Love et al., 2014). Gene ontology analysis was performed using Panther. The data files were submitted to GEO database: GSEXXXXXX codes will be available before publication.

### Immunohistology

Muscles were collected, immobilized on a piece of cork support using tragacanth gum and frozen in liquid nitrogen-cooled 2-methylbutane. Cross-sections 10 um thick were cut using a cryostat and transferred to Superfrost Plus Adhesion slides (Thermo Scientific). For each sample, three non-consecutive sections were processed for any given stain. For immunostaining, sections were equilibrated to room temperature, fixed with 4% paraformaldehyde and incubated for 1 hour with blocking solution (PBS with 10% normal serum). Primary antibodies were diluted in blocking buffer and incubation was performed overnight at 4C. Incubation with secondary antibodies, diluted in blocking buffer, was performed at room temperature for 1 hour. Fiber typing was performed with Development Studies Hybridoma Bank (DSHB, Iowa, USA) mouse monoclonal antibodies for MyHC-I (BA-D5), MyHC-IIa (2F7), MyHC-IIx (6H1) and MyHC-IIb (BF-F3). To facilitate visualization of fiber boundaries, sections were co-stained with Texas Red-X-conjugated wheat germ agglutinin (WGA). For visualization of blood vessels, we stained sections with anti-CD31 antibody (PECAM-1, R&D Systems AF3628) and WGA. Glut4 was detected using a rabbit polyclonal anti-Glut4 antibody (Sigma Aldrich 07-1404). All immunostains were mounted using Fluoroshield Media with DAPI (Sigma Aldrich) and imaged using a Zeiss LSM710 confocal microscope. Quantifications were performed using Fiji. To analyze NADH dehydrogenase activity we used NADH nitrotetrazolium blue stains (NADH-NTB). To that end, sections were incubated with freshly-prepared NADH-NTB solution (1.5 mM NADH, 1.5 mM NTB, 0.2 M Tris-HCl pH 7.4) for 15 minutes at 55C. Sections were washed with deionized water, dehydrated in graded ethanols, cleared in xylene, and mounted with Pertex mounting media (Histolab). For succinate dehydrogenase activity determination, sections were instead incubated in freshly-prepared 0.2 M phosphate buffer pH 7.6 supplemented with 170 mM succinic acid (Sigma-Aldrich) and 1.22 mM NTB (Sigma-Aldrich) for 30 minutes at 37C. For determination of fiber CSA, sections were stained with hematoxylin and eosin (H&E) and imaged using a green fluorescence filter. Imaging of NADH-TNB, SDH-NTB, and H&E stains was performed using a Zeiss AxioVert.A1 microscope. Quantifications were performed using Fiji.

### Immunoblotting

Frozen tissue was homogenized in protein lysis buffer (50 mM Tris-HCl pH 7.4, 180 mM NaCl, 1 mM EDTA, 1% Triton X-100, 15% glycerol) supplemented with protease and phosphatase inhibitor cocktails (Sigma-Aldrich). Cell cultures were washed with PBS and directly lysed in SDS lysis buffer (125 mM Tris-HCl pH6.8, 2% SDS, 10% glycerol) supplemented with protease and phosphatase inhibitor cocktails and sonicated in a Bioruptor (Diagenode). Protein lysates were quantified using the Pierce BCA Protein Assay Kit (Thermo Scientific) according to the manufacturer’s instructions. Protein lysates were resolved by SDS polyacrylamide gel electrophoresis and transferred to polyvinylidene membranes. Membranes were blocked with 5% skim milk and incubated with antibodies against NRTN (Thermo Scientific PA5-18769), SV2a (Abcam ab32942), MyHC-IIx (DSHB 6H1, MyHC-IIb (DSHB BF-F3), OXPHOS mitoprofile (Abcam ab110413), Glut4 (Sigma Aldrich 07-1404), Erk1/2 (CST 9101), p-Erk1/2 T202/Y204 (CST 9102), Creb (CST 9197), p-Creb S133 (CST 9198), Stat3 (CST 9139), p-Stat3 S727 (CST 9134), p-Stat3 Y705 (CST 9145), AKT (CST 9272), p-AKT S473 (CST 4058), S6 (CST 2217), p-S6 S235/236 (CST 4858), SRC (CST 2109), p-SRC family Y416 (CST 2101), or alpha-tubulin (Sigma-Aldrich T6199). Membranes were incubated with HRP-conjugated secondary antibodies and visualized by enhanced chemiluminescence on a ChemiDoc Imaging System (Bio-Rad).

### Human high-intensity sprint cycling

For analysis of *NRTN* gene expression in human muscle, we used vastus lateralis muscle biopsies from a previously published study (Schlittler et al., 2019). Muscle biopsies were obtained from healthy volunteers before and 72h after high-intensity sprint cycling.

### Publicly available datasets

Data for skeletal muscle Nrtn expression in different animal models were collected from the following studies deposited on GEO database: GSE23365 (PGC-1α mKO, PGC-1β mKO and PGC-1α+β mKO); GSE22086 (HSA-ERRγ); GSE40439 (MCK-PGC-1α1); GSE67049 (HSA-PGC-1α1 and HSA-PGC-1α-b); and GSE58699 (HSA-PGC-1β). Skeletal muscle Nrtn expression (Muscle mRNA; EPFL/LISP BXD CD+HFD Muscle Affy Mouse Gene 1.0 ST (Dec11) RMA) was correlated with BXD published phenotypes using the GeneNetwork database (www.genenetwork.org).

### Statistical Analysis

All statistical analysis were performed using GraphPad Prism Software. Quantitative data are presented as the mean +/-SEM. All parameters and number of samples are indicated in the corresponding figure legend. Unpaired Student’s t test was used to determine statistical significance when two groups were compared. For independent experiments with cultured cells, paired Student’s t test was used instead. One-way ANOVA followed by Fisher’s least significance difference (LSD) test for post hoc comparisons was used to determine statistical significance when multiple groups were compared. Statistical significance was defined as p-value <0.05 and denoted with asterisks.

## ACKNOWLEDGEMENTS

This study was supported by grants to J.L.R. and J.T.L. (Swedish Research Council 2016-00785, Novo Nordisk Foundation NNF16OC0020804, and the Swedish Diabetes Foundation DIA 2018-374) and to J.C.C. (The Lars Hiertas Memorial Foundation FO2014-0220, FO2015-0685, FO0216-0680). J.C.C. and

I. C. were partially supported by postdoctoral fellowships from the Swedish Society for Medical Research (SSMF) and P.R.J. received a postdoctoral fellowship from the Wenner-Gren Foundation.

## Author Contributions

J. C.C., Y.K., and J.L.R. conceived, coordinated and designed the study. J.C.C and Y.K. performed experiments with contribution from P.R.J., C.S., D.S., L.Z., I.C., M.O., J.N., V.M.R., M.S., and N.K. P.R.J. and I.C. performed bioinformatic analysis. J.C.C., Y.K. and L.Z. analyzed data. J.T.L., S.K., and E.H. designed experiments and edited the manuscript. J.C.C, Y.K. and J.L.R. wrote the manuscript.

## Declaration of Interests

S.K., M.S., and N.K. are employees and shareholders of Regeneron Pharmaceuticals, Inc. J.L.R. is a consultant for Bayer AG.

