## Supplementary Figs. for "Muscle-secreted neurturin couples myofiber oxidative metabolism and slow motor neuron identity"

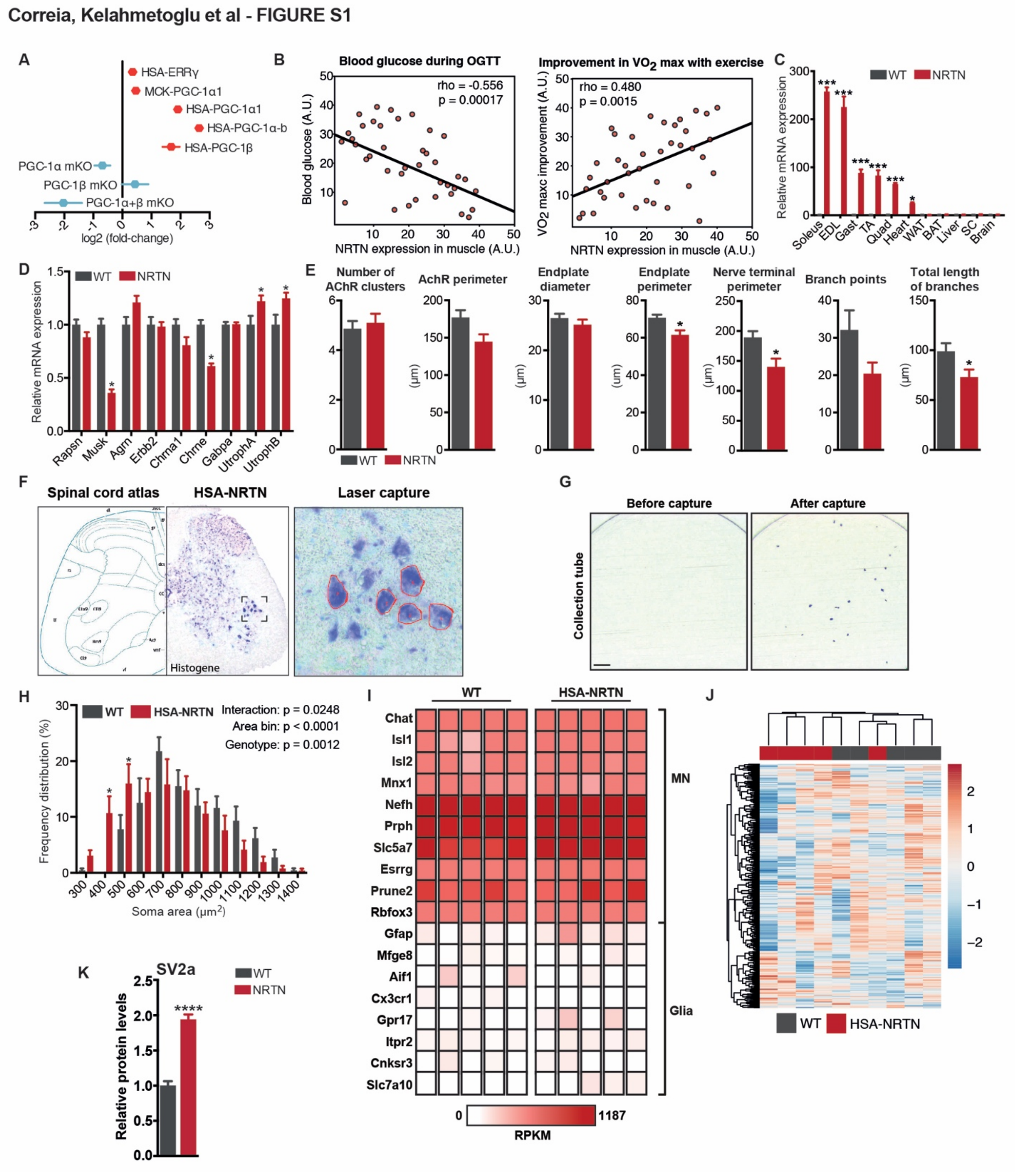


**Figure S1. NMJ and motor neuron remodeling in HSA-NRTN mice. Related to Figure 1.**

(A) *Nrtn* expression in skeletal muscle of PGC-1 transgenic and knockout mouse models.

(B) Correlations between skeletal muscle *Nrtn* expression and blood glucose during an oral glucose tolerance test and exercise-induced improvement in $VO_2$ max. Correlations were calculated using GeneNetwork based on transcriptomics and phenotypic data from BXD mouse strains.

(C) Analysis of *Nrtn* expression by qRT-PCR in multiple tissues from wild-type (WT) and HSA-NRTN mice (n = 4-6).

(D) Analysis of gene expression by qRT-PCR of gastrocnemius of WT and HSA-NRTN mice (n = 5-6).

**Figure S1 (continued).**

(E) Quantification of pre- and post-synaptic NMJ properties in WT and HSA-NRTN mice (n = 6). For a detailed flowchart of the NMJ morph analysis platform see (Jones et al., 2016).

(F) Spinal motor neurons visualized with Histogene staining were identified by location and morphology using the Spinal Cord Atlas as a reference.

(G) Laser captured motor neurons collected on a tube cap for further processing for RNA-seq.

(H) Histogram showing the size distribution of WT and HSA-NRTN spinal motor neurons collected by laser-capture microdissection.

(I) Heatmap showing the expression levels (RPKM) of motor neuron and glial markers in cells captured from the lumbar spinal cord of WT and HSA-NRTN mice.

(J) Heatmap summary of changes in gene expression in motor neurons from HSA-NRTN mice, compared to those from WT mice (n = 5).

Bars depict mean values and error bars represent SEM. * $p<0.05$, ** $p<0.01$, *** $p<0.001$, **** $p<0.0001$.

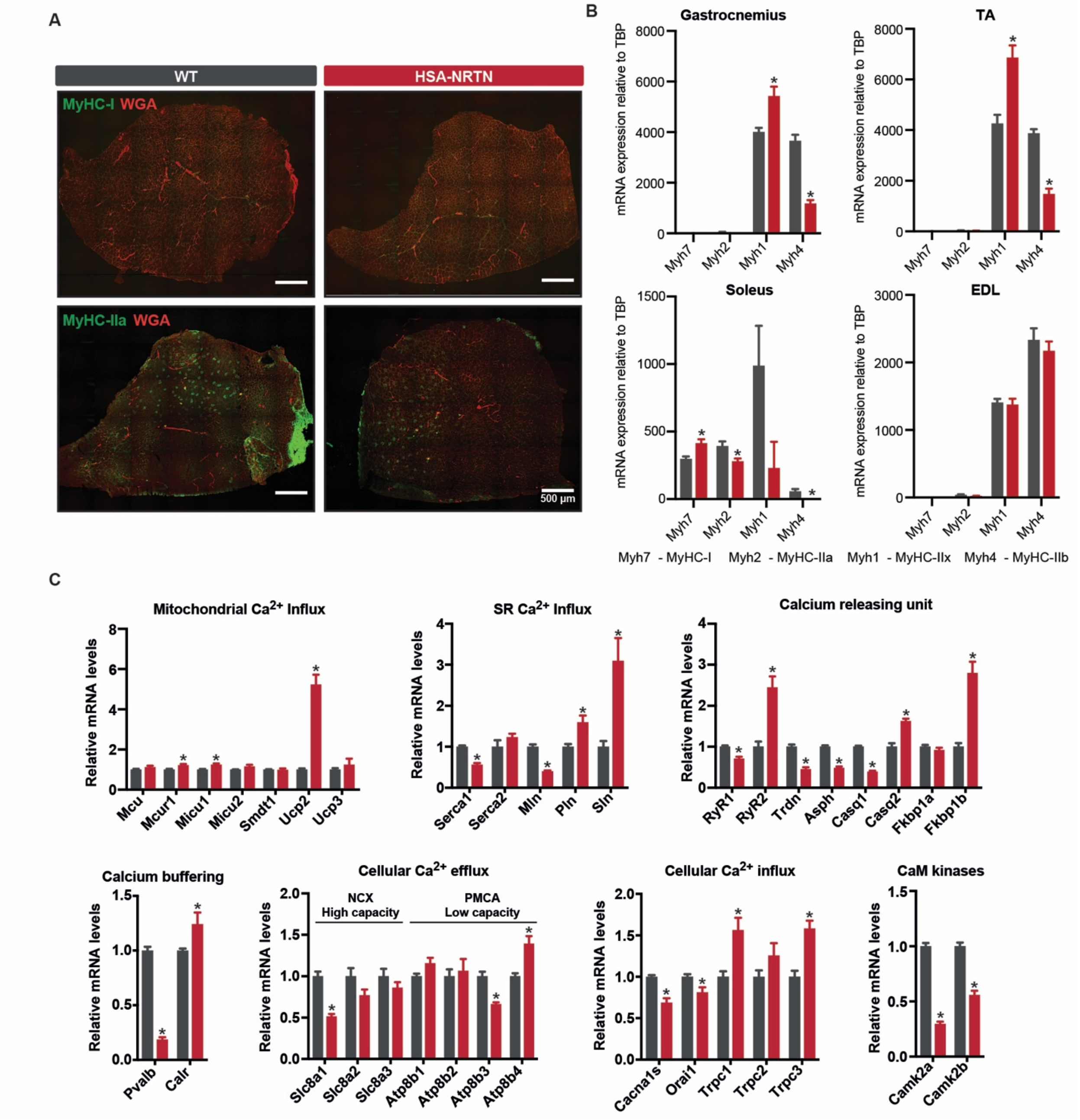


**Figure S2. Transcriptional reprogramming of contractile and calcium handling machinery in HSA-NRTN skeletal muscle. Related to Figure 2.**

(A) Immunohistochemical analysis of fiber type composition in TA muscles of WT and HSA-NRTN mice. Cell membranes are visualized using wheat germ agglutinin (WGA, in red).

(B) Gene expression analysis by qRT-PCR of myosin heavy chain isoforms in gastrocnemius, TA, soleus, and EDL muscles from HSA-NRTN and wild-type (WT) littermates (n = 5-8)

(C) Gene expression analysis of calcium handling machinery by qRT-PCR in gastrocnemius muscles from HSA-NRTN and WT mice (n = 6).

Bars depict mean values and error bars represent SEM. * $p<0.05$.

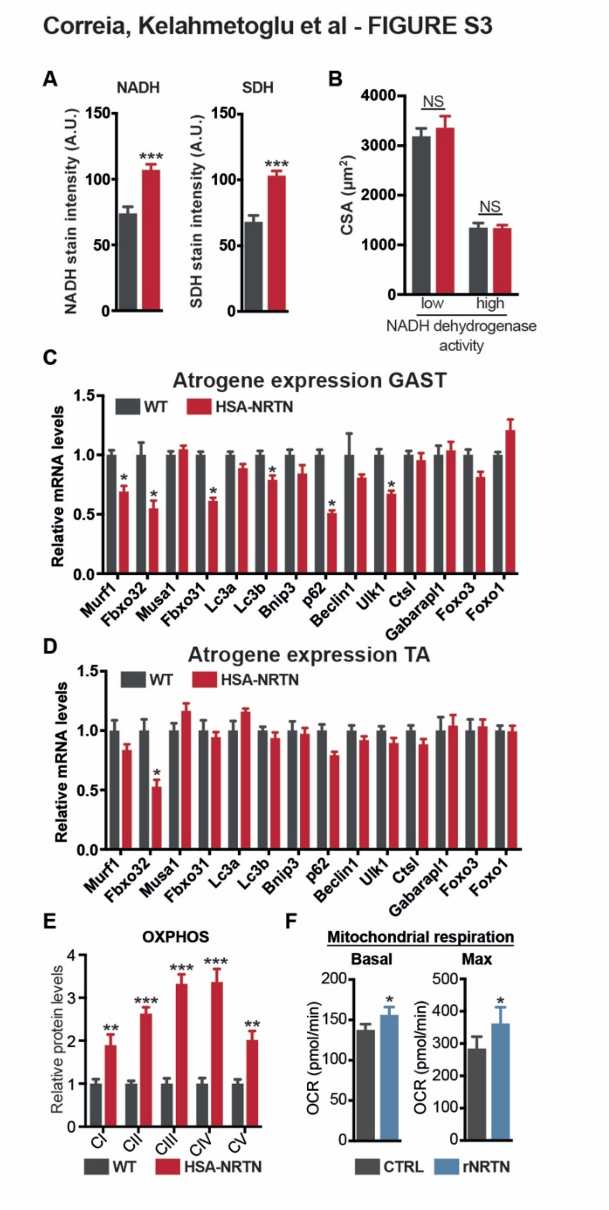


**Figure S3. NRTN does not activate an atrophy gene program in skeletal muscle. Related to Figure 3.**

(A) Quantification of NADH dehydrogenase and succinate dehydrogenase activity stains in TA muscles HSA-NRTN mice, compared to wild-type (WT) mice (n = 5-7).

(B) Quantification of fiber cross-sectional area (CSA) in TA fibers with high or low NADH dehydrogenase activity (n = 5-7).

(C-D) Gene expression analysis by qRT-PCR of atrogenes in gastrocnemius (C) and TA (D) muscles from HSA-NRTN mice, compared to WT controls (n = 6).

(E) Quantification of immunoblots for components of the mitochondrial electron transport chain in gastrocnemius muscles of HSA-NRTN mice, compared to WT controls (n = 4).

(F) Quantification of basal and maximal mitochondrial respiration in mouse primary myotubes treated with 10 ng/ml rNRTN for 24h (n = 3).

Bars depict mean values and error bars represent SEM. * p<0.05, ** p<0.01, *** p<0.001.

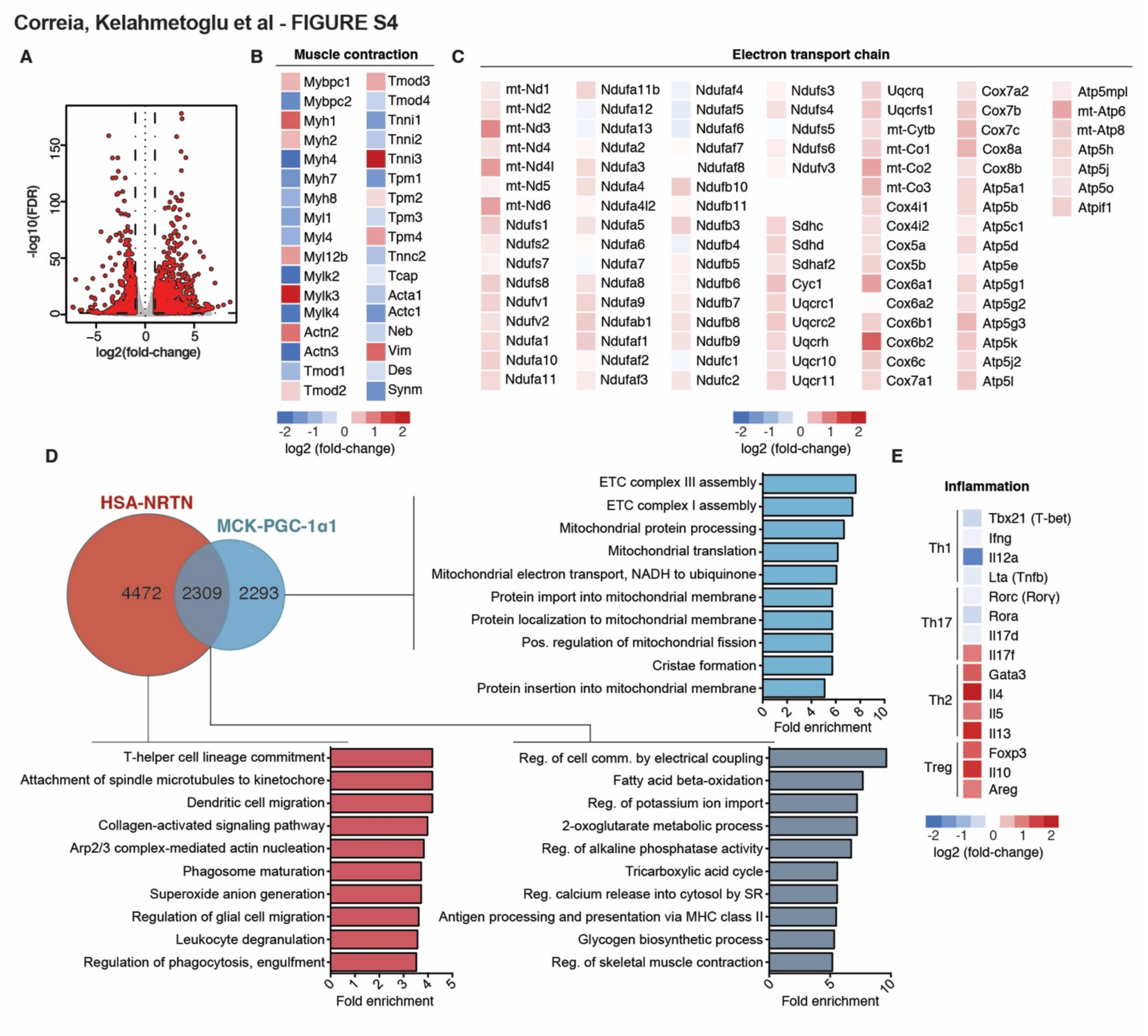


**Figure S4. RNA-sequencing analysis of HSA-NRTN skeletal muscle. Related to Figure 4.**

(A) Volcano plot of differentially expressed genes in HSA-NRTN gastrocnemius, compared to wild-type (WT) controls (n = 3).

(B-C) Heatmaps showing the relative changes in expression of genes encoding components of the muscle contractile machinery (B) and mitochondrial electron transport chain (C) (n = 3).

(D) Comparison of differentially expressed gene sets from HSA-NRTN and MCK-PGC-1α1 skeletal muscle. The Venn diagram illustrates the overlap between both gene sets. Gene ontology analysis was performed for the overlapping gene set and for each specific gene set.

(E) Heatmaps showing the relative changes in expression of markers of T cell populations (n = 3).

Correia, Kelahmetoglu et al - FIGURE S5

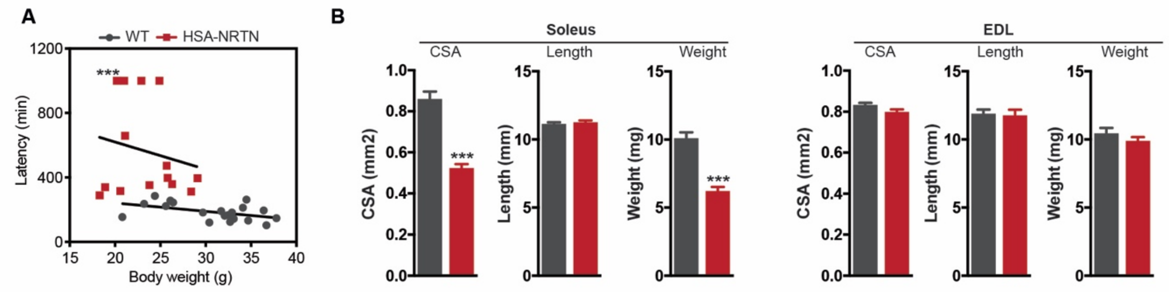


**Figure S5. NRTN improves motor coordination and exercise performance. Related to Figure 5.**

(A) Correlation between body weight and rotarod performance (day 3) of wild-type (WT) and HSA-NRTN mice.

(B) CSA, length and weight of soleus and EDL muscles from WT and HSA-NRTN mice (n = 4).

Bars depict mean values and error bars represent SEM. *** $p<0.001$.

Correia, Kelahmetoglu et al - FIGURE S6

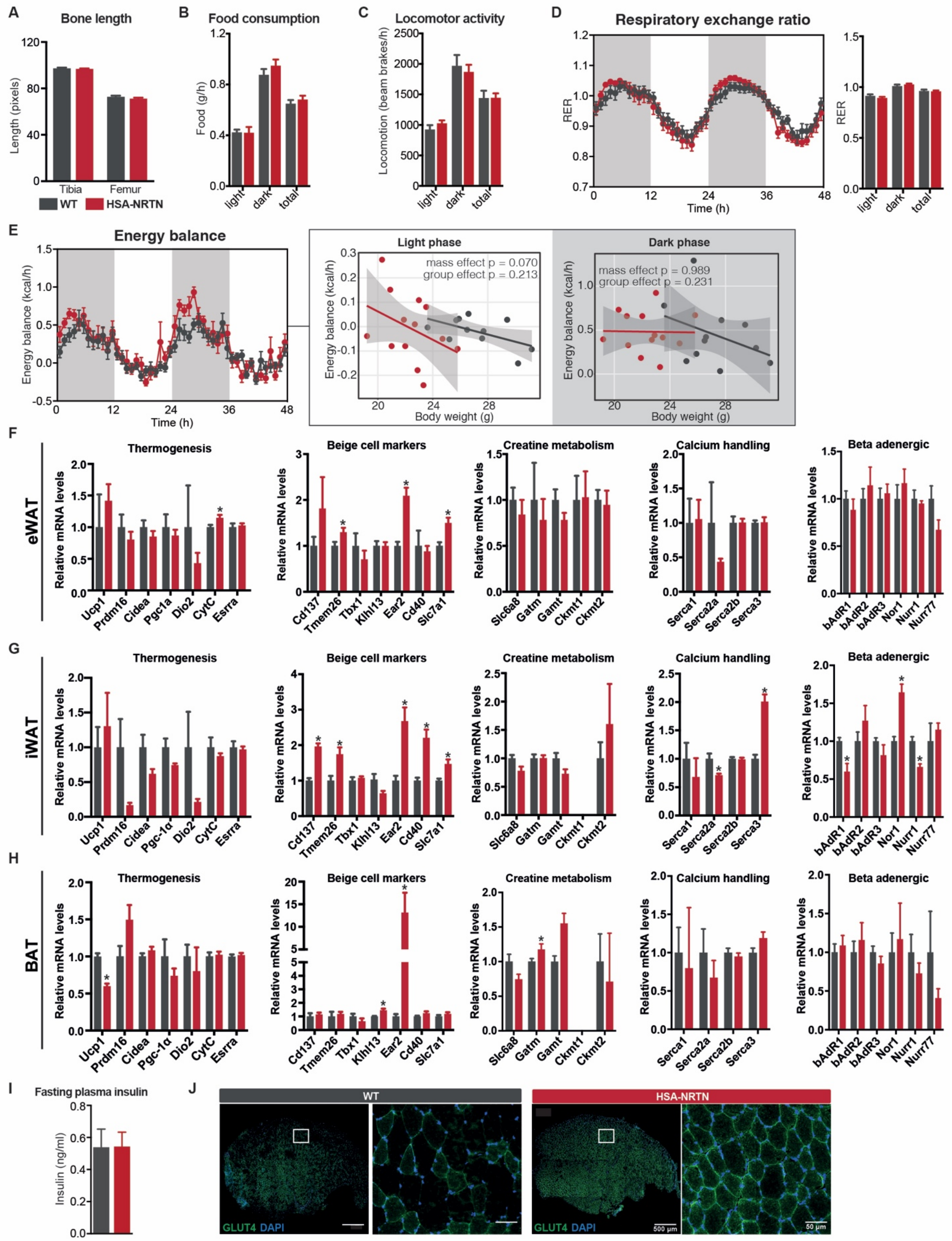


**Figure S6. Metabolic characterization of HSA-NRTN mice. Related to Figure 6.**

(A) Tibia and femur length measured from DEXA scan images of wild-type (WT) and HSA-NRTN mice (n = 8).

**Figure S6 (continued)**

(B-E) Food consumption (B), locomotor activity (C), respiratory exchange ratio (D), and energy balance (E) determined by CLAMS in WT and HSA-NRTN mice (n = 8).

(F-H) Gene expression analysis by qRT-PCR of thermogenic gene programs in eWAT (F), iWAT (G), and BAT (H) of HSA-NRTN mice, compared to WT controls (n = 5-6).

(I) Fasting (5h) plasma insulin levels in WT and HSA-NRTN mice (n = 3-5).

Bars depict mean values and error bars represent SEM. * $p<0.05$.

(J) Immunostaining for GLUT4 in TA muscles of WT and HSA-NRTN mice.

Correia, Kelahmetoglu et al - FIGURE 7

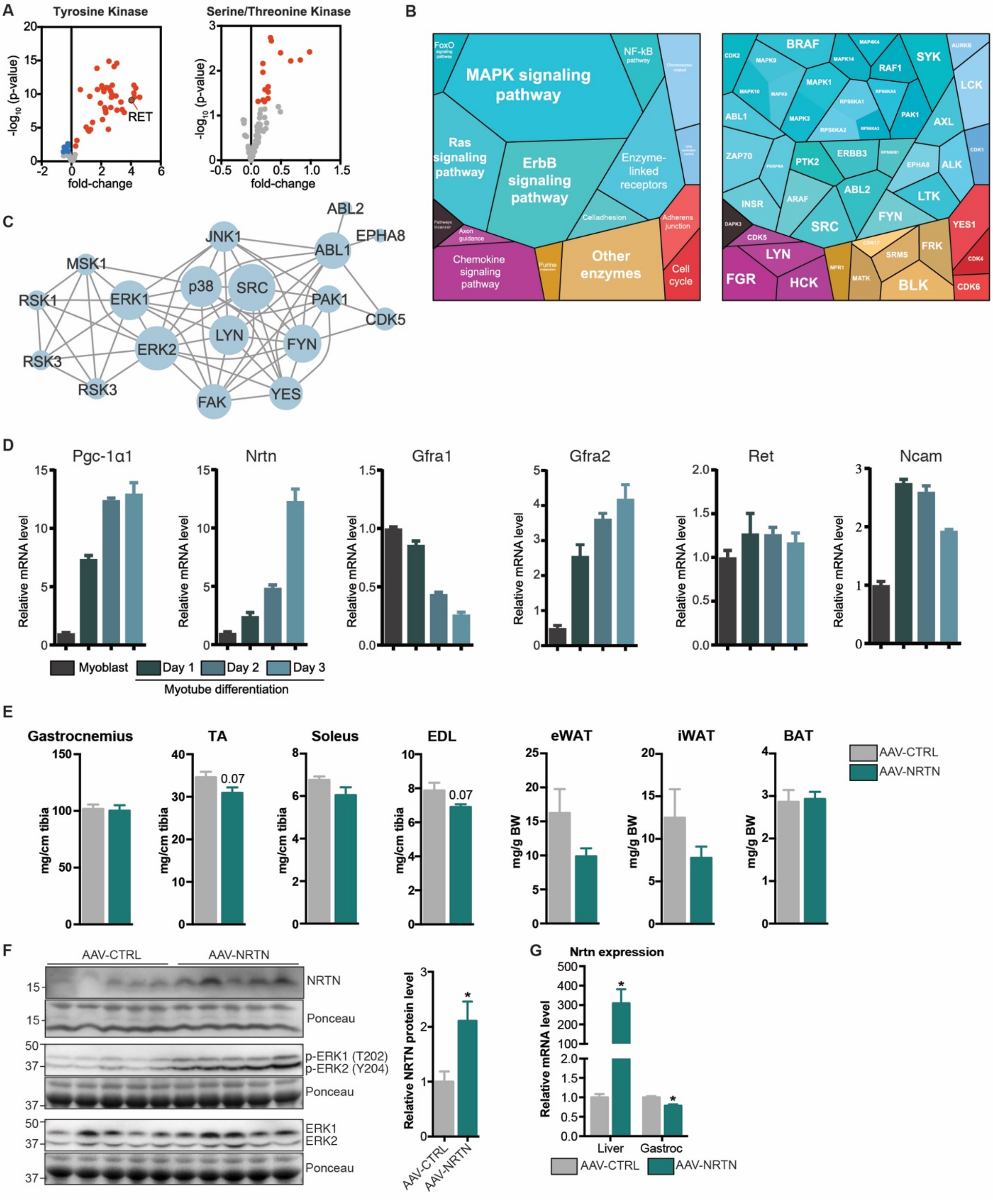

**Figure S7. Kinase activity profiling in HSA-NRTN skeletal muscle. Related to Figure 7.**

(A) Relative changes in peptide phosphorylation in HSA-NRTN gastrocnemius muscles, compared to wild-type (WT) controls (n = 3-5).

(B) Proteomaps depicting differentially active kinases and associated pathways in HSA-NRTN gastrocnemius muscles, compared to WT controls.

(C) Axon guidance protein-protein interaction subnetwork, generated using STRING and Cytoscape from differentially active kinases that are part of the axon guidance pathway.

**Figure S7 (continued).**

(D) Gene expression analysis by qRT-PCR of mouse primary myotubes at different stages of differentiation. Data from a single experiment performed in triplicate.

(E) Tissue weights of C57BL/6N mice injected intravenously AAV8 vectors encoding a construct expressing *Nrtn* (AAV8-NRTN) or a control construct (AAV8-CTRL) (n=5).

(F) Immunoblot analysis of NRTN and ERK1/2 phosphorylation in gastrocnemius muscles of mice injected with AAV8-CTRL or AAV8-NRTN (n = 5).

(G) *Nrtn* gene expression in liver and gastrocnemius muscle of mice injected with AAV8-CTRL or AAV8-NRTN (n = 5).

Bars depict mean values and error bars represent SEM. * p<0.05.
